# Multiplexed smFISH reveals the spatial organization of neuropil localized mRNAs is linked to abundance

**DOI:** 10.1101/2024.07.13.603387

**Authors:** Renesa Tarannum, Grace Mun, Fatima Quddos, Sharon A. Swanger, Oswald Steward, Shannon Farris

## Abstract

RNA localization to neuronal axons and dendrites provides spatiotemporal control over gene expression to support synapse function. Neuronal messenger RNAs (mRNAs) localize as ribonucleoprotein particles (RNPs), commonly known as RNA granules, the composition of which influences when and where proteins are made. Highthroughput sequencing has revealed thousands of mRNAs that localize to the hippocampal neuropil. Whether these mRNAs are spatially organized into common RNA granules or distributed as independent mRNAs for proper delivery to synapses is debated. Here, using highly multiplexed single molecule fluorescence in situ hybridization (HiPlex smFISH) and colocalization analyses, we investigate the subcellular spatial distribution of 15 synaptic neuropil localized mRNAs in the male and female rodent hippocampus. We observed that these mRNAs are present in the neuropil as heterogeneously sized fluorescent puncta with spatial colocalization patterns that generally scale by neuropil mRNA abundance. Indeed, differentially expressed mRNAs across cell types displayed colocalization patterns that scaled by abundance, as did simulations that reproduce cell-specific differences in abundance. Thus, the probability of these mRNAs colocalizing in the neuropil is best explained by stochastic interactions based on abundance, which places constraints on the mechanisms mediating efficient transport to synapses.

**SIGNIFICANCE STATEMENT:** RNA localization establishes compartment-specific gene expression that is critical for synapse function. Thousands of mRNAs localize to the hippocampal synaptic neuropil, however, whether mRNAs are spatially organized as similar or distinctly composed ribonucleoprotein particles for delivery to synapses is unknown. Using multiplexed smFISH to assess the spatial organization of 15 neuropil localized mRNAs, we find that these mRNAs are present in variably sized puncta suggestive of heterogeneous transcript copy number states. RNA colocalization analyses in multiple hippocampal cell types suggest that the spatial relationship of these mRNAs is best predicted by their abundance in the neuropil. Stochastic RNA-RNA interactions based on neuropil abundance are consistent with models indicating that global principles, such as energy minimization, influence population localization strategies.

## INTRODUCTION

Neuronal morphology is incredibly complex, and in order for neurons to function efficiently messenger RNA (mRNA) transcripts need to be delivered to distant sites for on-demand translation. In particular, mRNA localization to synapses and subsequent local translation are required for the synaptic plasticity underlying learning (Bradshaw et al., 2003; Huber et al., 2000). Dysregulation of these processes is a common cause of intellectual disability and autism (Fernandopulle et al., 2021; Huber et al., 2002). Thus, uncovering how mRNA cargoes are delivered to and locally regulated at the synapse is central to our understanding of the molecular basis of learning and memory. Furthermore, studying the fundamental principles of neuronal mRNA localization can uncover key aspects of post-transcriptional regulation, which could be applicable to various other organisms and cell types, such as yeasts, drosophila germ cells, cardiomyocytes, etc., that use compartmentalization for gene regulation (Lewis et al., 2018; Martin & Ephrussi, 2009).

The hippocampus is a brain region critical for learning and memory that has a laminar organization ideal for cataloging and visualizing localized mRNAs in the axon- and dendrite-rich neuropil layer. Compartment-specific deep sequencing studies have revealed the presence of thousands of mRNA species localized in the rodent hippocampal synaptic neuropil (Cajigas et al., 2012; Farris et al., 2019). These localized mRNAs are assembled and transported throughout the neuropil as “ribonucleoprotein particles” (RNPs, otherwise known as RNA granules), which are dynamic, spherical, membraneless macromolecular complexes of nontranslating mRNAs, mRNA-binding proteins (RBPs), and translational machinery (Kiebler & Bassell, 2006; Knowles et al., 1996). Several flavors of RNPs localize to the synaptic neuropil, including transport RNPs, RISC RNPs, translating RNPs, p-bodies, and stress granules (Bauer et al., 2023; Kiebler & Bauer, 2024). These RNPs are typically characterized by their RBP constituents (e.g., FMRP (Antar et al., 2005), DCP1, AGO2 (Cougot et al., 2008; Zeitelhofer et al., 2008), CPEB (Huang et al., 2003), EIF4E (Napoli et al., 2008), G3BP (Sahoo et al., 2018), staufen (Sharangdhar et al., 2017) despite significant heterogeneity in composition due to the overlap and exchange of RBPs between RNP types. Recent in vitro studies in neurons and human cell lines show that mRNA is a key driver of mRNA-protein condensates by influencing RNP composition and size through seeding of higher order molecular assemblies (Bauer et al., 2022; Garcia-Jove Navarro et al., 2019; Maharana et al., 2018). However, investigations into the mRNA composition of RNPs and whether it contributes to molecular specificity have traditionally been overlooked.

Several different models have been proposed to explain how mRNAs are assembled into RNPs to dictate their destination. One hypothesis is that mRNAs are transported as single or low copy number molecules per RNP. Single-molecule fluorescence in situ hybridization (smFISH) studies in cultured neurons revealed that individual dendritic RNAs, whether in transport or localized, carry no more than one or two molecules of a specific type of transcript (Batish et al., 2012; Mikl et al., 2011). Similar studies also revealed whether specific pairs of dendritically localized mRNAs co-exist in common particles (Gao et al., 2008) or not (Batish et al., 2012; Mikl et al., 2011; Tübing et al., 2010). However, larger scale observations were precluded due to limitations in multicolor RNA labeling. Thus, there is limited evidence on the heterogeneity (how much of a given mRNA) and diversity (how many types of different mRNAs) of neuronal RNP compositions as well as their spatial distribution in intact neural circuits. Nevertheless, selective delivery of low copy number RNPs appears at odds with sustaining the localization of thousands of diverse mRNAs in the synaptic neuropil with varying abundances and encoded protein functions (Ainsley et al., 2014; Cajigas et al., 2012; Farris et al., 2019). In contrast, immunoprecipitation studies from brain lysates or dissociated cultured neurons suggest that mRNAs selectively associate within larger mRNA granules that contain many different types of transcripts and RBPs, some that are selective for specific granules (Elvira et al., 2006; Fritzsche et al., 2013; Heraud-Farlow et al., 2013; Kanai et al., 2004). Although this model seems plausible and efficient for localizing vast amounts of mRNAs, these studies are technically limited by the lack of spatial resolution and non-specific RNA interactions. Addressing this question requires subcellular-resolution imaging of many endogenous molecules at once, which is now technically feasible using highly multiplexed, single molecule fluorescence in situ hybridization with iterative imaging (HiPlex smFISH).

In this paper, we used single, 3-color, and HiPlex smFISH to characterize the spatial distributions of localized neuronal RNAs in the rodent hippocampal neuropil. We generally focused on neuropil localized mRNAs that are targets of FMRP (fragile X messenger ribonucleoprotein), a ubiquitous RBP involved in neuronal mRNA localization and translational regulation (Richter et al., 2015). FMRP is associated with nearly 400 localized mRNAs in the hippocampal neuropil (Hale et al., 2021; Sawicka et al., 2019). However, it remains unclear whether localized FMRP mRNA targets segregate into distinct or similarly composed RNPs, which could contribute to the diversity and/or selectivity of FMRP-RNP compositions. Here we show, based on heterogeneity in mRNA fluorescent puncta area, that 15 neuropil localized RNAs likely vary in individual transcript copy numbers, existing as either low or high copy number populations, or more frequently, as both populations within the same mRNA. Simultaneous visualization of 12 neuropil localized FMRP-target mRNAs revealed that these mRNAs are spatially organized as such that their pairwise co-distribution, assessed as colocalization, is mostly explained by their abundance in the neuropil. When assessing the colocalization of all 12 RNAs at once, the highly abundant mRNAs were overwhelmingly present in mRNA clusters defined by the presence of three or more mRNAs. This remained true when mRNA clusters were defined by the presence of FMRP protein. We further show that cell-specific differences in colocalization can largely be explained by differences in abundance. Lastly, we used simulations to show that increased mRNA abundance can achieve the colocalization levels observed in experimental images. Collectively, these data provide evidence that, for the hippocampal mRNAs studied here, mRNAs are localized to the neuropil in heterogeneously sized puncta that may reflect differences in individual mRNA copy number and display colocalization patterns that are best explained by neuropil abundance. These data from intact rodent hippocampal neuropil are generally in agreement with imaging studies from cultured neurons, suggesting that both systems are subject to similar intrinsic mechanisms that favor independent localization with stochastic overlaps as opposed to coordinated assembly of these RNAs into selective multimeric granules in the neuropil.

## MATERIALS & METHODS

### Animals

Sexually naive adult female Sprague Dawley rats were used for the *Arc* dilution studies. Both male and female C57BL/6J were used at p17 for the HiPlex experiment and 8-16 weeks of age for Shank2 and 3plex studies. Animals were group-housed under a 12:12 light/dark cycle with access to food and water ad libitum. All procedures were approved by the Animal Care and Use Committee of Virginia Tech or University of California Irvine and were in accordance with the National Institutes of Health guidelines for care and use of animals.

### Stimulation Paradigm

The stimulation paradigm was as previously described (Steward and Worley, 2001) with the following modifications. Briefly, an electroconvulsive seizure (ECS) was induced in unanesthetized adult female Sprague Dawley rats by delivering AC current (60Hz, 40mA for 0.5s). Anesthesia was induced immediately after ECS by intraperitoneal injection of 20% urethane. The animals were then placed in a stereotaxic apparatus and a stimulating electrode was positioned to selectively activate one side of the medial perforant path projections (1.0 mm anterior to transverse sinus and 4.0 mm lateral from the midline). The electrode depth was empirically determined to obtain a maximal evoked response in the dentate gyrus at a minimal stimulus intensity, typically 3-4mm deep from the cortical surface. A recording electrode was positioned in the molecular layer of the dorsal blade of the dentate gyrus (3.5 mm posterior from bregma, 1.8 mm lateral from the midline, 3-3.5 mm from the cortical surface based on evoked responses generated by stimulation). Single test pulses were then delivered at a rate of 1/10 s for 20 minutes to determine baseline response amplitude. Two hours after the ECS delivery, high frequency stimulation (trains of eight pulses at 400hz) were delivered at a rate of 1/10 s. After 60 minutes the brain was removed and flash frozen. Brains were embedded in OCT and sectioned in the coronal plane on a cryostat at 20 μm and processed for FISH as described below.

### Fluorescence in situ hybridization (FISH)

FISH was performed as previously described (Guzowski et al., 1999; Farris et al 2014) to examine *Arc* mRNA puncta in dendritic fields of the dentate gyrus. For the dilution experiments, a 1X saturating stock of full-length dig-labeled *Arc* probe was serially diluted with full length unlabeled *Arc* probe at 1:2, 1:4, and 1:8.

### Quantitative Analyses of *Arc* puncta number and size

Sections processed for FISH were imaged across the molecular layer of the dentate gyrus at 63X using a confocal microscope. The size and number of *Arc*-positive puncta were determined using imageJ particle analysis function (NIH). Briefly, the images were overlaid using DAPI and a region of interest (ROI) was determined so as to count each *Arc*-puncta only once. The entire image was then set to a threshold calculated as the average automated otsu threshold across undiluted images, which was confirmed to show little to no detectable signal on negative control slides, and then the thresholded images were watershed to segment individual *Arc*-puncta. The ROI was then cropped out of the original image and subjected to particle analysis. The number and feret’s diameter of *Arc*-puncta at each dilution were averaged across three sections (technical replicates) from one animal and data are presented as mean +/- SEM across sections (technical replicates).

### Single Molecule Fluorescence in situ hybridization (smFISH)

Brains were embedded in OCT and sectioned in the horizontal plane (mouse studies) on a cryostat at 20 μm and processed for smFISH according to the RNAscope Fluorescent Multiplex or HiPlex kit instructions (Advanced Cell Diagnostics, Hayward, CA). RNAscope in situ hybridization probes can efficiently detect single mRNA transcripts (F. Wang et al., 2012) and smFISH RNA signals detected by this commercially available kit are strongly correlated with RNA sequencing read counts from dissected hippocampal neuropil (Farris et al., 2019). The following mus musculus specific probes were used with the RNAscope fluorescent multiplex reagent kit (Cat# 320850): *Rgs14* (Cat #416651), *Adcy1* (Cat #451241), *Ppp1r9b* (Cat #546311), *Shank2*-O2 Pan (Cat #513711, NM_001113373.3/ENSMUST00000105900.8), *Shank2*-O3 Short 2a (Cat # 851661-C2, ENSMUST00000146006.2/NM_001113373.3), *Shank2*-O4 long 2e (Cat # 852961-C3, ENSMUST00000105900.8/NM_001081370.2), mouse 3 plex positive control (Cat # 320881), 3 plex negative control (Cat # 320871). Mus musculus specific probes used for HiPlex assay (Cat# 324400) includes *Adcy1* (Cat #451241-T1), *Aco2* (Cat #1120581-T2), *Psd* (Cat #449711-T3), *Dlg4* (Cat #462311-T4), *Calm1* (Cat #500461-T5), *Bsn* (Cat #1119681-T6), *Camk2a* (Cat #445231-T7), *Pum2* (Cat #546751-T8), *Ddn* (Cat #546261-T9), *Pld3* (Cat #507241-T10), *Ppfia3* (Cat #1119691-T11), *Cyfip2* (Cat #561471-T12), HiPlex positive control (Cat #324321) and negative control (Cat #324341).

### *Shank2* smFISH, image acquisition, and analysis

*Shank2* smFISH was performed according to the instructions provided in the kit (Cat# 320851). Probes labeling *Shank2e*-long, *Shank2a*-short and *Shank2-*pan were imaged with Alexa-647, Atto-555 and Atto-488, respectively. ROIs captured from CA2 cell body were 211 µm X 211 µm in x-y plane and 5 µm in z (25 steps, step size: 0.21 µm) at 63X magnification using a Leica thunder (Leica DMi 8) widefield epifluorescence microscope. Images were denoised and deconvolved in NIS elements AR (v5.41.01) using Richardson-Lucy deconvolution algorithm to increase signal to noise ratio and remove background. After all computational processing steps for signal optimization, maximum projection images were used for further analysis. A binary segmentation layer was created using the “bright spot” command in NIS AR based on the fluorescent intensity of the smFISH puncta. Segmentation was based on intensity threshold chosen according to the corresponding negative control image. For quantification of colocalization (defined as >1% overlap) between mRNA puncta in individual channels, union binary layers were created using the “having” command, e.g. “Shank2a puncta having Shank2-pan puncta”, to measure the number of overlapping puncta. Colocalization values were calculated as the percentage of overlapping puncta relative to the total number of puncta for that individual channel of interest. Example equation is as follows:

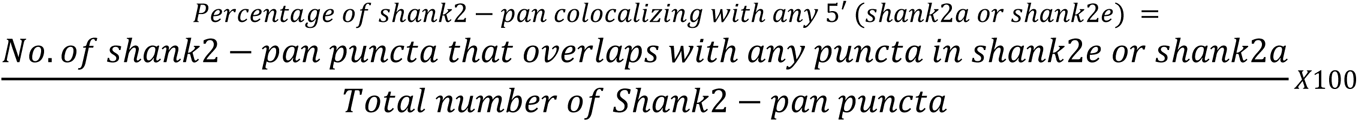

To calculate the percentage of puncta that would be randomly colocalized (Dunn et al., 2011; McDonald & Dunn, 2013; Sauerbeck et al., 2020), one image from each pair was rotated 180 degrees due to the diagonal orientation of the CA2 cell body layer in the acquired images and then the colocalization values were repeated and presented as ‘random’. Paired one-tailed t-tests were performed due to a priori expectation that the experimental colocalization would be greater than the random overlap. It is worth mentioning that colocalization in fluorescence microscopy can be quantified using pixel-based correlation coefficients or object-based segmentation methods (Bolte & Cordelières, 2006; Dunn et al., 2011). Global intensity-based correlation coefficients, such as Pearson’s and Manders’, are susceptible to background noise and uneven illumination, and can only detect relationships between two channels of interest. Since our objective was to assess colocalization—i.e., the spatial relationship between at least two and up to twelve different smFISH signals detected in separate channels—we opted for an object-based approach. The distinct architecture and margins of discrete fluorescent spots in high-resolution multiplex and HiPlex smFISH images provided an advantage, enabling effective segmentation of signals for subsequent object-based colocalization metrics.

### HiPlex smFISH

HiPlex smFISH was performed on slide mounted 20 µm sections using RNAscope HiPlex Assay V2 (Cat# 324400). After fixation, samples were dehydrated in % ethanol and treated with protease IV for 30 mins. Samples were hybridized at 40℃ with the twelve probes for 2 hours followed by signal amplification steps. T1-T4 fluorophores were added to label *Adcy1, Aco2, Psd* and *Dlg4* mRNAs in round one. For each round, 488, 550, 647 and 750 nm LED were used to image four mRNAs at 63X magnification (Numerical aperture 1.4). Leica Thunder epifluorescence microscope (Leica DMi 8) was used for imaging with recorded stage positions to acquire the same ROIs across rounds. Individual channel acquisition parameters were selected to optimize signal per mRNA in the experimental images with less than 1-3% of the number of experimental puncta in the associated negative control images. *Adcy1* mRNA label was in round one as a CA2 marker and DAPI signal was acquired using a 390 nm LED. After round one, coverslips were taken off by keeping slides in 4X SSC, fluorophores were cleaved and FFPE reagent was used to decrease background. Subsequently T5-T8 fluorophores were added to image *Calm1, Bsn, Camk2a* and *Pum2* in the second round. This was followed by similar steps of cleaving the fluorophores and background removal. For the final round, T9-T12 fluorophores were added to image *Ddn, Pld3, Ppfia3, Cyfip2* mRNAs. Exposure was adjusted in each round matching with the expression of individual mRNAs but kept consistent across all animals per run (N=2 mice per run for 2 separate runs). After completion of smFISH round 3, fluorophores were cleaved and slides were washed in TBS for 2X5 mins, blocked in TSA-blocking solution for 30 mins and incubated with anti-rabbit-FMRP primary antibody (1:100, Abcam, Cat# ab17722, Lot# 632949982) at 4C for two consecutive nights. Subsequently slides were washed in TBS-T (0.05% Tween) for 3X5 mins and 2% H_2_O_2_ in TBS for 10 mins at room temperature. Following that, slides were incubated with goat-anti-rabbit HRP (1:250, Jackson Immunoresearch, Cat# 111035144, Lot# 149770) for 2 hours at room temperature. Slides were washed in TBS-T before they underwent incubation with TSA-Cy3 (1:50, Cat# NEL704A001KT, Lot# 210322048) for 30 mins at room temperature. Slides were then washed in TBS-T for several times and coverslipped with prolong gold antifade mounting medium (refractive index 1.51). Images of FMRP immunostaining were done using tissue from N=2 animals, imaged using 550 LED and signals were adjusted using the (no primary) negative control slide. ROIs from proximal and distal neuropil of CA1 and CA2 were imaged that were 211 µm X 211 µm in x-y plane and 5 µm in z-plane (step size 0.21 µm) at 63X magnification.

### HiPlex smFISH image analysis

All z-stack images of individual channels and rounds were exported as TIFF images and converted to nd2 format for further processing on NIS elements AR. Denoising ai and Richardson-Lucy deconvolution algorithm was used to increase the signal to noise ratio and minimize background pixel intensity. Images were then maximum projected and registered using ACD RNAscope HiPlex image registration software (version 1.0.0) based on the DAPI signal of each round. After registration, a composite image of 12 mRNA channels and a DAPI channel (plus the FMRP channel for N=2 mice) was created for segmentation. Experimental images and negative control images were processed identically at acquisition and post-processing. Each channel was segmented to a binary layer, using intensity threshold, based on the fluorescence intensity of the experimental image and corresponding negative control channel (bacterial gene *DapB* with T1-T12 channel specific fluorophores) such that the negative control produced little to no segmented objects (<3% of detected puncta of the experimental image). Any segmented fluorescent puncta overlapping DAPI + 10% surrounding area was removed from the further analysis to examine only neuropil localized mRNAs. Binaries were optimized manually for every channel to best represent the ground truth mRNA signal. Fluorescent puncta area data was exported per mRNA channel for N=4 mice and then plotted as relative percentage area distribution which was subsequently averaged for plotting on the heatmap.

For the colocalization analysis, four 52 X 52 µm^2^ ROIs were cropped from the 211 X 211 µm^2^ image. For each mRNA, subsequent binary layers were created that would contain only mRNA puncta from the channel of interest having any overlap from each of the other channels/ mRNAs in consideration. Thus, eleven binaries were created for each mRNA channel to calculate the number of that mRNA having overlapped puncta with any of the other eleven mRNAs. This number was then expressed as a percentage of the given mRNA.

For the quantification of the random level of overlaps, the same method of calculation was followed only after rotating the image of the given mRNA to 90 degrees right as well as 180 degrees. Both 90 degrees and 180 degrees rotated images resulted in a similar percentage of random overlap so only data from 90-degree rotated images are shown. Percentage of random colocalization was subtracted from the percentage of colocalization calculated from experimental images and plotted in a heatmap. Calculation of *Psd* % colocalization with any mRNA in HiPlex dataset was done by creating a union layer of all intersect binary layers for *Psd*. Thus “*Psd* having *Camk2a* (mRNA 1)”, “*Psd* having *Ddn* (mRNA 2)”…..”*Psd* having *Ppfia3* (mRNA 11)” all layers were merged to create a union layer that includes *Psd* puncta that has overlapping objects from any of the other 11 channels. Similarly, correlation of mRNA colocalization with mRNA abundance was calculated for *Ddn* and *Pum2*. For subsequent analysis, additional binary layers were created per channel including only mRNA signals that colocalized with FMRP and pairwise colocalization and *Psd* mRNA composition analysis were repeated similarly on this subset.

### Rgs14, Adcy1 and Ppp1r9b smFISH

*Rgs14*, *Adcy1*, and *Ppp1r9b* smFISH was performed using RNAscope fluorescent multiplex reagent kit (#Cat 320851) as described in the kit protocol. 211 µm X 211 µm ROI (in x-y plane) of 5 µm thickness Z-stack images (25 steps, step size 0.21µm) were acquired by a Leica thunder (Leica DMi 8) wide-field fluorescence microscope at 63X magnification (Numerical aperture 1.4). CA2 was identified using *Rgs14* and *Adcy1* labeling. *Rgs14, Adcy1* and *Ppp1r9b* were imaged using 488, 550 and 647 nm LEDs. CA1 and CA2 proximal and distal neuropil regions and DG were imaged from N=4 adult mouse hippocampus. Exported TIFF images were then processed using NIS elements AR. Each image including negative control was computationally processed by denoising and Rich-Lucy deconvolution algorithm. A binary segmentation layer was created, using intensity threshold and based on the fluorescent intensity of the experimental and corresponding negative control image, per channel on the post-processed max projection images and manually edited to best represent the data. Any mRNA puncta overlapping plus 10% of DAPI was excluded from quantification to confirm only mRNAs in the neuropil but not in glia or interneurons are included in the analysis. After manually editing each binary layer, the number of mRNA puncta and fluorescent puncta area data was exported to excel and plotted with prism.

### Rgs14, Adcy1 and Ppp1r9b smFISH image analysis

180 µm X 180 µm^2^ ROI was cropped from CA2, CA1 and DG images for object-based colocalization analysis and colocalization between individual channels was defined as minimally touching (1%) to 100% overlapping binary objects from two separate channels of interest. The number of mRNA puncta that colocalized between two channels was then divided by the total number of mRNA puncta in both channels and plotted as a percentage.

For simulation of CA2 images, object-based segmentation allowed us to export XY coordinates, puncta area, intensity and ferret’s diameter of all neuropil mRNA puncta in each channel (including interneuron clusters) from chosen ROIs (180 X 180 µm^2^) in CA2 proximal dendrite images and DG (N=4 mice). These images were segmented similarly as above for 3Plex mRNAs except without removal of puncta overlapping DAPI signal to avoid bias in the specific XY coordinates or puncta area in randomly simulated data (all the previous analysis in this paper included mRNA puncta only in the neuropil not overlapping DAPI to exclude interneuron specific signals). *Adcy1* mRNA puncta simulation analysis was done in python (3.9.13) in spyder (5.2.2). The difference between the number of *Adcy1* mRNA puncta between CA2 vs DG from the same brain section was calculated by subtraction and then a list of equal number of random *Adcy1* mRNA puncta were created using exported CA2 *Adcy1* mRNA puncta properties (highest and lowest centroid XY coordinates, feret’s diameter, area, sum intensity) using pandas, NumPy, and scipy.spatial packages (Virtanen et al., 2020). Colocalization of *Adcy1*/ *Ppp1r9b* mRNA was then recalculated in CA2 simulated images to compare with experimental DG images (>1% overlap was defined if the distance between centroid XY coordinates of two puncta is less than or equal to 0.99 X sum of radius (feret’s diameter/2) of each puncta pair). Ten iterations were done for one CA2 proximal dendrite image per mouse and then the average of colocalized *Adcy1/Ppp1r9b* mRNAs was calculated per mouse for N=4 mice. For quantification of >50% overlap data a similar pipeline was used (>50% overlap was defined if the distance between centroid XY coordinates of two puncta is less than or equal to 0.50 X sum of radius (feret’s diameter/2) of each puncta pair).

Fluorescent puncta area data of *Rgs14, Adcy1, Ppp1r9b* mRNAs from proximal and distal neuropil of CA2 and CA1 and entire molecular layer of DG (211 µm X 211 µm) was first plotted in prism as individual histograms of each animal (data not shown). Although the total number of mRNA puncta varied for each mRNA species from mouse to mouse, a consistent pattern of area distribution was noted when plotted as a percentage fraction. Due to consistent patterns across neuropil laminae, data from proximal and distal neuropil were then combined to represent the size histogram as the relative percentage of mRNAs of different sizes for CA2, CA1 and DG was plotted as a heatmap. Bin width was kept consistent for all RNA puncta area distribution plots. For description of the data, area data were averaged for each cell type across mice and then averaged across cell types.

### Statistical analyses

All statistical tests were paired-two-tailed except for comparison of experimental and random colocalization which were one-tailed due to the a priori hypothesis that experimental colocalization would be higher than randomly colocalized puncta. All statistical analyses were done using Graphpad Prism (v10) with a significance level of 0.05 or lower (□=0.05).

## RESULTS

### *Arc* mRNAs contain multiple copies of *Arc* transcripts

To begin to address whether neuronal mRNAs are localized in the neuropil as low- and/or multiple copy number-containing RNA puncta, we investigated the RNA properties of the well-known neuropil localized mRNA *Arc* (activity-regulated cytoskeleton-associated mRNA). *Arc* mRNA expression in the hippocampal dentate granule (DG) cell dendrites is unique in that it is tightly regulated by activity-dependent transcription and degradation (Farris et al., 2014). At baseline, dendritic *Arc* expression is low or absent in most DG cells, but after a single electroconvulsive shock (ECS) *Arc* mRNA is rapidly transcribed (within ∼3 minutes) and transported throughout the DG dendritic laminae by 30 min to 1 hour (Steward et al., 1998). Given the short half-life of *Arc* mRNA (∼45 min (Rao et al., 2006), the prolonged presence of *Arc* mRNA in DG dendrites (e.g. at 2 hrs.) is maintained by ongoing transcription and dendritic transport (Farris et al., 2014). Subsequent unilateral high frequency stimulation (HFS) of the entorhinal cortical perforant path inputs to the DG further boosts *Arc* transcription and leads to the accumulation of newly transcribed *Arc* mRNA selectively near the activated synapses in the middle molecular layer and a depletion of *Arc* mRNA from the outer molecular layer (Steward et al 1998; Farris et al 2014). Using this stimulation paradigm (ECS + HFS) on a single adult female rat and fluorescence in situ hybridization (FISH), we assessed the size and number of dendritically-localized (ECS) and synaptically-localized (HFS) *Arc* mRNA puncta to examine whether *Arc* mRNA composition changes with synaptic localization (Fig. 1). We found that *Arc* RNAs are generally of similar size, as measured by feret’s diameter, whether they are localized to dendrites or targeted to recently activated synapses (Fig. 1AB, average from three technical replicates). RNAs are generally larger in distal versus proximal dendrites under both conditions. These data suggest that resident and newly transcribed dendritic *Arc* RNAs contain a similar amount of *Arc* mRNA per puncta.

**Fig. 1:**
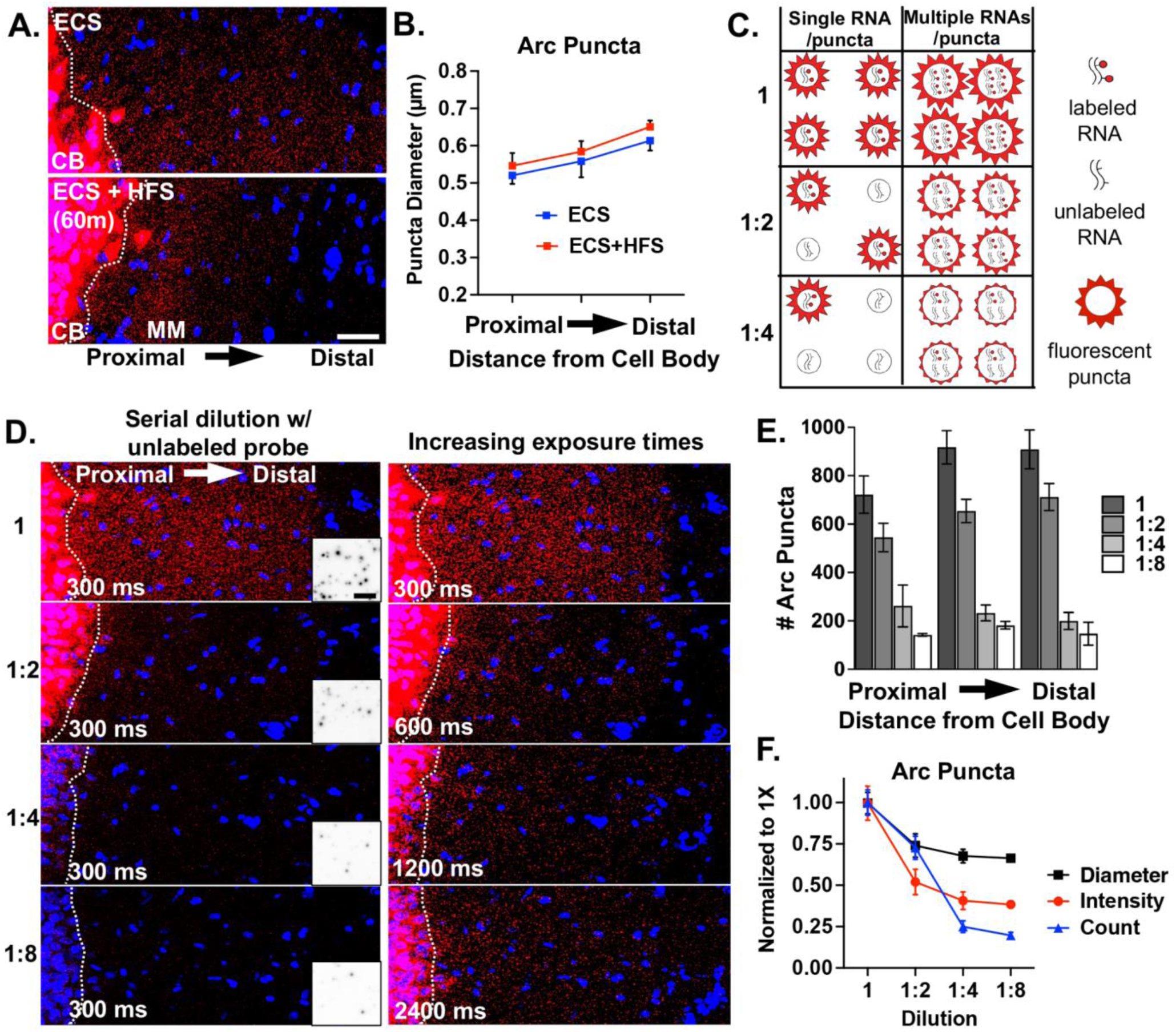
Arc mRNA fluorescent puncta diameter, number, and intensity upon probe dilution reveal multiple pools of RNAs. **A.** Representative images of *Arc* mRNA localization to the middle molecular layer (MM) of the rat dentate gyrus following ECS and 60 min unilateral HFS. Scale = 25 µm. **B.** Quantification of the Feret’s diameter of localized *Arc* mRNA puncta (HFS condition) vs. non-localized puncta (ECS) by distance from the cell body layer (CB, white dashed lines). Average diameter ± SEM per layer from n=3 technical replicates. Approximately 500-1000 puncta per replicate as in E (1X dilution). **C.** Possible outcomes of serial probe dilution on single and multiple copy number *Arc* puncta in terms of number and intensity or apparent size of fluorescent puncta. **D.** Representative images after ECS only labeled with 1X undiluted full length *Arc* probe or serially diluted with unlabeled full length *Arc* probe and imaged with identical exposure times, 300 ms (Left). Images acquired with doubling exposure times (600, 1200, 2400 ms) revealed undetected puncta at 300 ms (Right), indicating a decrease in puncta intensity as would be expected with multiple RNAs per puncta. Inverted inset scale = 2.5 µm. **E.** Quantification of *Arc* puncta number for each dilution at 300 ms. Stepwise decrease in *Arc* puncta number suggests low copy number containing puncta. Average number ± SEM per layer from n=3 technical replicates. **F.** Quantification of *Arc* puncta Feret’s diameter, intensity, number for the middle molecular layer of each dilution at 300 ms. Normalized average ± SEM from n=3 technical replicates. A decrease in diameter and/or intensity with dilution suggests a pool of multiple copy number containing puncta. See representative inverted inset images in D.

Next, in order to assess the transcript occupancy of *Arc* puncta, we measured fluorescent puncta diameter and number in rats that received an ECS only (dendritically localized) after serial dilution of 1X labeled full length *Arc* probe with unlabeled (cold) full length *Arc* probe (1:2, 1:4, 1:8). We reasoned that a stepwise decrease in *Arc* RNA puncta number would reflect mRNAs transported singly or at low copy numbers that were no longer detectable when half, fourth, or eighth of the probe was labeled (Fig. 1C). Alternatively, a decrease in apparent fluorescent puncta size would reflect a reduction in the number of labeled transcripts from multiple copy-containing *Arc* RNA puncta (Fig. 1C). When acquiring images using identical acquisition parameters (optimized for 1X labeled probe), we detected a stepwise decrease in *Arc* RNA puncta number, with the largest drop off between one half and one fourth cold probe dilution (Fig. 1DE). These data are consistent with a population of low copy number *Arc* RNAs. However, when we doubled the exposure time after each dilution, we qualitatively saw an increase in the number of *Arc* RNA puncta indicating that a proportion of the *Arc* RNAs labeled with cold probe dropped below the detection threshold (Fig. 1D). This is in agreement with the measured decrease in fluorescence intensity and apparent size (feret’s diameter) of *Arc* RNA punctas between undiluted and diluted conditions (Fig. 1F), which we interpret to reflect a population of *Arc* RNA puncta containing multiple copies of *Arc* transcripts. Collectively, these data suggest that there are multiple populations of *Arc* RNAs, those with both low and high *Arc* copy numbers that exist in DG dendrites. Given that we did not detect any changes in RNA puncta size after HFS, we assume this finding would translate to synaptically targeted *Arc* RNAs.

### smFISH probes can detect mRNA colocalization

In addition to neuropil localized RNAs consisting of low or multiple copies of the same mRNA (homotypic), we wanted to test whether they are composed of multiple species of mRNAs (heterotypic) as described in situ for established RNA granules, like germ plasm granules (Trcek et al., 2015) or p-bodies (Cougot et al., 2008; Ford et al., 2019) and identified biochemically for neuronal transport granules (Elvira et al., 2006; Fatimy et al., 2016; Fritzsche et al., 2013; Heraud-Farlow et al., 2013). In order to assess the colocalization of different mRNA transcripts into common RNA puncta, we first needed to confirm that we can reliably detect colocalized smFISH signals. To test this, we took advantage of the fact that there are (at least) two isoforms of *Shank2* expressed in hippocampal area CA2 (Farris et al 2019). The two isoforms are generated via alternative 5’ promoters and thus differ in their 5’ untranslated regions (UTRs), but have identical 3’ UTRs (Jiang & Ehlers, 2013; Monteiro & Feng, 2017). Using isoform specific probes targeted to the two distinct 5’ UTRs (Shank2e-long and Shank2a-short) and a pan Shank2 probe targeted to the common 3’UTR (Shank2-pan, Supp. Fig. 1A), we calculated the percentage of colocalized signals, defined as puncta in separate channels overlapping by at least 1%. In agreement with RNAseq expression data (Supp. Fig. 1A), we detected *Shank2* expression from all three probes in area CA2 (Supp. Fig. 1BC). In general, the Shank2-pan probe detected more *Shank2* mRNAs than the 5’ UTR probes combined (# of RNAs: Shank2-pan = 3056 ± 472, Shank2e = 1122 ± 268, Shank2a = 1463 ± 79, N= 3 mice), either due to the (presumed) greater accessibility of the 3’ UTR from less RNA secondary structure compared to the 5’ UTRs, or potentially due to expression of other isoforms that include the 3’UTR but not either of the two 5’UTRs (e.g. Shank2C, (Monteiro & Feng, 2017)) that cannot be resolved via short-read sequencing. We found that nearly 40% of Shank2e (35.41 ± 2.57%) or Shank2a (41.53 ± 7.05%) colocalized with the Shank2-pan probe (Supp. Fig. 1D). The Shank2-pan probe also colocalizes with either 5’ UTR probe at ∼30% (29.88 ± 2.42%). The fact that this relative percentage is not greater than the colocalization of the individual 5’ UTR colocalization is due to both the greater number of Shank2-pan labeling (described above) and several instances where all three probes colocalized at presumed transcriptional foci (Supp. Fig. 1B, inset). These data are consistent with our previous findings, where ∼30% of 5’ and 3’ Arc mRNA probes colocalize in the dendrites of dentate granule cells in rats (Farris et al., 2014). Thus, we reason that RNAscope smFISH is more limited in its ability to detect colabeling of the same individual RNA transcript with two probes (∼30%), perhaps due to steric hindrance or competition of the DNA-based labeling approach, but it is highly likely to detect colocalization when more than one transcript is being labeled (e.g. two transcripts of the same RNA or two distinct neuropil localized RNAs).

To account for the amount of colocalization expected to occur by randomly overlapping puncta, which is also influenced by expression levels, we rotated one of the channels from each probe pair 180 degrees and remeasured “random” colocalization (Dunn et al., 2011; McDonald & Dunn, 2013). Image rotation is a validated technique for assessing random colocalization of synaptic molecules in the hippocampal neuropil (Frye et al., 2021; Sauerbeck et al., 2020). Here, because the cell body layer is in a diagonal orientation, we rotated 180 degrees instead of the commonly used 90 degrees. In most cases, we detected a significantly greater % colocalization than was observed at random (Supp. Fig. 1D). In the instance where % colocalization is near random, as with the two 5’ probes, we assume this to indicate that these two transcripts do not colocalize often into common complexes. In summary, our method is able to reliably detect colocalization of two probes targeted to the same mRNA, which is a higher bar than for detecting two mRNAs within the same RNA puncta, which we assess below.

### Putative FMRP-target mRNAs have heterogeneous puncta area distributions

Subcellular localization of a given mRNA is assumed to be affected by the composition of the RNA-RBP complexes (Mikl et al., 2011; Mitsumori et al., 2017). Once we demonstrated that our method can reliably detect colocalization when we expect it, we explored whether known neuropil mRNA transcripts localize independently or in association (colocalized) with each other as heterotypic complexes of two or more RNAs. We rationalized that mRNAs with a shared RBP interactor would be more likely to demonstrate colocalization patterns reflecting some degree of selectivity in how they associate with each other, if at all. We took advantage of the relatively well characterized RBP, FMRP, and its neuropil localized target mRNAs to quantitatively map their colocalization at subcellular resolution in the mouse hippocampus. We generated a list of candidate target mRNAs by cross-referencing datasets that identified hundreds of putative FMRP target mRNAs using HITS-CLIP (high-throughput sequencing of mRNA isolated by crosslinking immunoprecipitation) on whole brain (Darnell et al., 2011) and hippocampal CA1 neuropil (Sawicka et al., 2019) with datasets that identified high-confidence hippocampal neuropil mRNAs (Ainsley et al., 2014; Cajigas et al., 2012; Farris et al., 2019). This list of neuropil localized candidate FMRP target mRNAs was further curated based on expression, different encoded protein functions (signaling, cytoskeletal, synaptic plasticity, etc.) and target destinations (mitochondria, cytoplasm, cell membrane, dendritic spine) to further stratify colocalization patterns (Supp. Table 1).

To spatially map the association of these putative FMRP target mRNAs, we probed for 12 endogenous mRNAs at once and iteratively imaged 4 at a time using HiPlex smFISH followed by FMRP immunostaining (Supp. Fig. 2). Experimental and negative control images were post-processed with denoising and deconvolution and each mRNA channel was segmented based on an intensity threshold that produced negligible signal in the corresponding negative control image (Supp. Fig 3). Each of the mRNAs were present in CA2 dendrites with varying degrees of abundance (Supp. Fig. 4). Based on the findings from our previous experiments (Fig. 1), which indicated the presence of distinctly sized populations of homotypic RNA puncta within the same mRNA species, we first calculated the median fluorescent puncta area (Supp. Table 2) and plotted the relative percent size distributions (Fig. 2AB). Unexpectedly, we found these mRNAs vary considerably in their fluorescent puncta area distributions, which are not explained by fluorophore, imaging round, or abundance in the neuropil (Supp. Table 1).

**Fig. 2.**
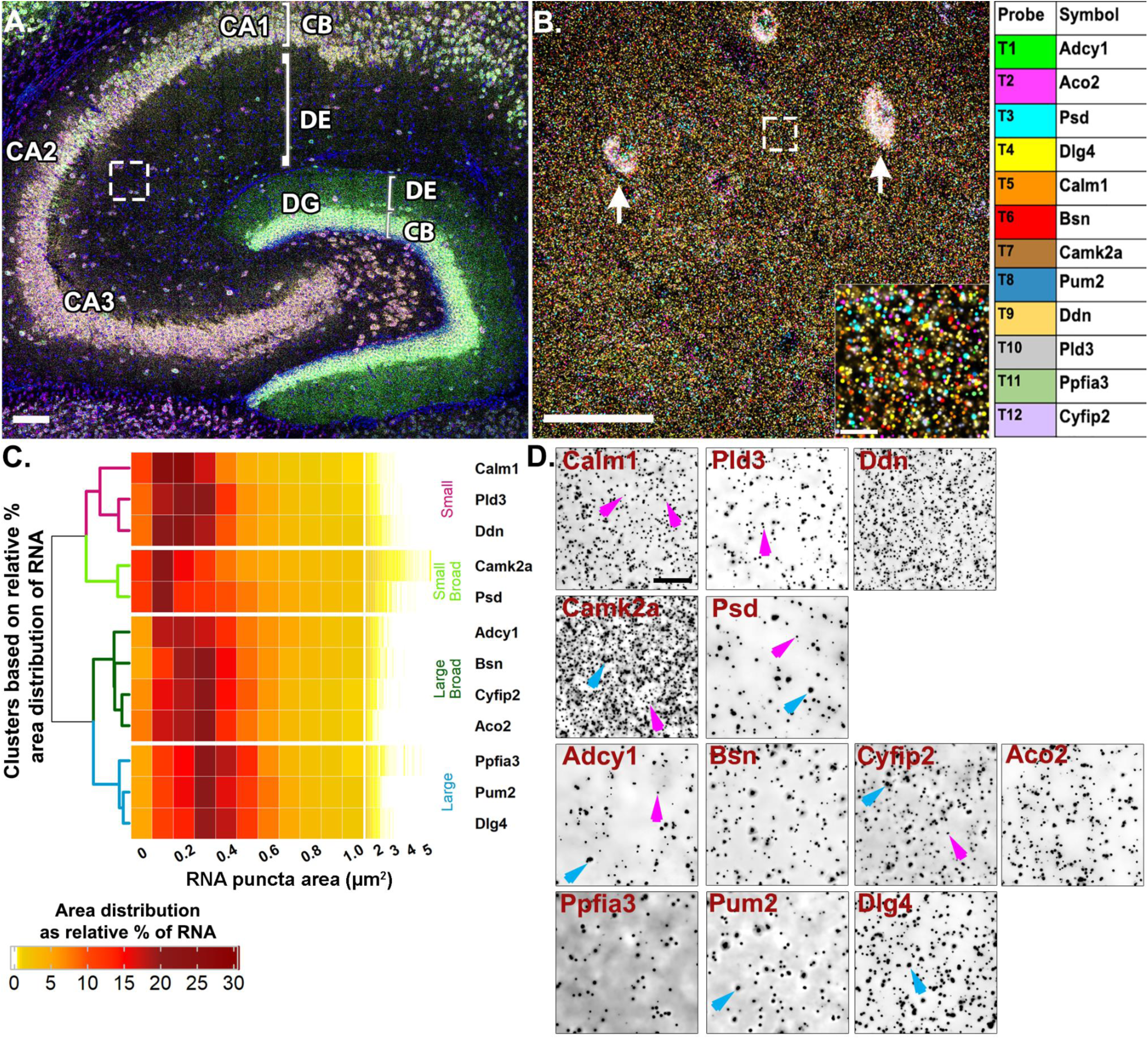
Highly multiplexed mRNA imaging reveals neuropil localized mRNAs have distinct puncta area distributions. **A.** Representative image of mouse hippocampus with *Adcy1, Aco2, Psd, Dlg4* labeling in round 1 of HiPlex smFISH. White box represents ROI from CA2 dendrites. **B.** Representative high-magnification merged image of 12 mRNAs from CA2 dendrites. Arrows denote inter-neuron expression in the neuropil layer that is removed before analysis (see methods) and inset is the dashed white box. Each mRNA is colored based on the table on the right. **C.** Heatmap of RNA fluorescent puncta area distributions hierarchically clustered by similarity. (N = 4 mice) **D.** Representative inverted images of each mRNA. Magenta arrows denote small RNAs in *Calm1*, *Pld3*, *Camk2a*, *Psd*, *Adcy1* and *Cyfip2*. Blue arrows denote larger sized RNAs in *Camk2a*, *Adcy1*, *Cyfip2*, *Ppfia3* and *Dlg4*. Scale = A. 100 µm, B. 50 µm, 10 µm, D. 10 µm.

Unsupervised hierarchical clustering analysis of the puncta area distributions revealed four different patterns (Fig. 2CD, Supp. Table 1). The first cluster is comprised of mRNAs with consistently “small” RNAs (*Ddn*, *Pld3* and *Calm1*, labeled magenta in the dendrogram, Fig. 2C), whereby, on average, ∼55% of RNAs are less than 0.3 µm^2^ (54.57 ± 4.84%, averaged across mRNAs from N=4 mice) with the largest relative percent peak (23.69 ± 1.77%) at 0.2 µm^2^. These consistently small RNAs have fewer than 10% RNAs (9.29 ± 1.46%) sized 0.6-1.0 µm^2^ and only 3% (2.50 ± 0.64%) of RNAs larger than 1.0 µm^2^. In contrast, the last cluster is comprised of mRNAs with consistently “large” RNAs (*Dlg4*, *Pum2* and *Ppfia3*, labeled blue, Fig. 2C) whereby, on average, ∼50% of the RNAs are 0.3-0.5 µm^2^ (49.80 ± 1.16%) with the largest relative peak (20.33 ± 0.42%) at 0.3 µm^2^. These consistently large RNAs have less than 5% RNAs larger than 1.0 µm^2^ (3.14 ± 0.26%). There are two intermediary clusters, “small broad” (*Camk2a* and *Psd*, labeled light green, Fig. 2C) and “large broad” (*Adcy1*, *Bsn*, *Aco2* and *Cyfip2*, labeled dark green, Fig. 2C) that have relatively broader size distributions that segregate with either the “small” or “large clusters”, respectively. The “small broad” cluster (*Camk2a* and *Psd*) shows a very broad distribution with the largest relative peak (21.21 ± 0.72%) equal to or less than 0.2 µm^2^ and a larger population of RNAs sized 0.6-1.0 µm^2^ that accounts for more than 15% (15.89 ± 1.71%). These “small broad” mRNAs have the largest fraction of RNAs greater than 1.0 µm^2^ at nearly 10% (9.38 ± 1.34%). The “large broad” cluster (*Adcy1*, *Bsn*, *Aco2* and *Cyfip2*) also shows a broad distribution with the largest relative peak (38.85 ± 1.12%) between 0.2-0.3 µm^2^ and a larger population of RNAs sized 0.6-1.0 µm^2^ that accounts for ∼15% (15.31 ± 1.05%). However, these “large broad” RNAs have only ∼3% of RNAs larger than 1.0 µm^2^ (2.69 ± 0.60%). Small and large populations are denoted on the representative images with magenta and blue arrows, respectively (Fig. 2D).

It is interesting to note that even some of the most abundant neuropil mRNAs visualized here contain populations with consistently small RNA puncta areas (i.e., *Ddn*, *Calm1*). These data suggest that mRNAs, regardless of abundance, vary considerably in RNA puncta area, both within a transcript population and across different transcripts. Consistent with the *Arc* probe dilution results, we interpret larger RNA puncta areas to likely represent RNA complexes with multiple copies of the same transcript, whereas the smaller RNAs likely represent RNA complexes containing fewer copies of transcripts or perhaps a single copy.

### Putative FMRP-target mRNAs colocalize in the neuropil based on abundance

To systematically characterize whether any particular FMRP-target mRNAs display similar colocalization profiles (and thus suggestive of co-regulation), we measured the number of overlapping fluorescent puncta (>1% overlap) between two mRNA channels to determine pairwise colocalization values (Batish et al., 2012; Carson et al., 2008; Gao et al., 2008). For each pair across the 12 mRNAs, we expressed the colocalization values as a percentage of each individual mRNA (Fig. 3, Supp. Fig. 5) and as a percentage of the combined pair (Supp. Fig. 6). We included well characterized neuropil localized RNAs (*Camk2a*, *Dlg4* also known as Psd95, *Cyfip2, Ddn*) and uncharacterized mRNAs (*Aco2*, *Psd*, *Pld3*). The degree of colocalization across pairs in properly registered experimental images ranged from 4.21 ± 0.48% (*Adcy1/Ppfia3*) to 71.38 ± 4.91% (*Camk2a*/*Ppfia3*) (Supp. Fig. 5A). The degree of colocalization that was observed at random (one image from every pair rotated 90 degrees, which controls for the differences in expression across mRNA pairs) ranged from 3.07 ± 0.41% (*Aco2*/*Ppfia3*) to 53.74 ± 4.75% (*Pum2*/*Camk2a*) (Supp. Fig. 5B). We then subtracted the random colocalization percentage from the percentage obtained from the properly registered experimental images, anticipating that random colocalization subtraction would eliminate the relationship with abundance, and visualized the result as a heatmap (Fig. 3A). The range of colocalization percentages above random spanned from 0.53 ± 0.36% (*Adcy1*/*Ppfia3*) to 27.74 ± 5.70% (*Psd*/*Camk2a*). Thus, after correcting for random colocalization, some mRNAs were rarely colocalized whereas others showed ∼20-30 times more colocalization, suggesting a difference in the propensity of mRNA species to be colocalized.

**Fig. 3.**
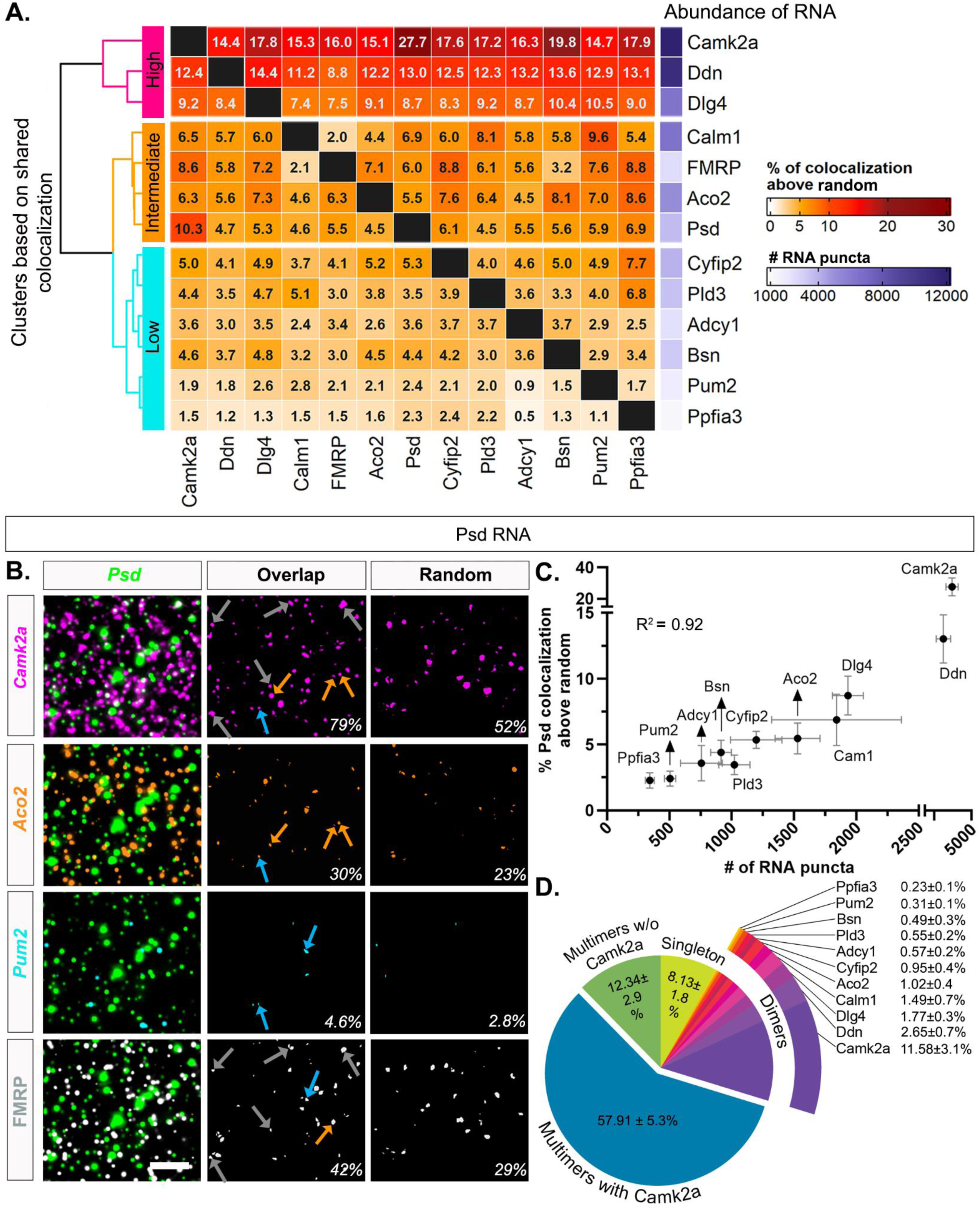
mRNA pairwise colocalization patterns correlate with abundance. **A.** Percentage of RNAs colocalized in pairwise combinations above random. Percentage was calculated by dividing the number of overlapping puncta by the total number of RNAs that correspond to each channel (columns), then subtracting the percentage obtained after rotating one of the images 90 degrees (See Supp. Fig. 5). **B.** Representative images of colocalization with *Psd*. The first column consists of merged images of *Psd* (green,) with i) *Camk2a* (magenta), ii) *Aco2* (orange), iii) *Pum2* (cyan) and iv) FMRP protein (white). The middle column shows the intersecting area of each pair of RNAs *Psd*/*Camk2a* (magenta, 85 out of 107 or 79% of *Psd* RNA colocalize with *Camk2a* in this image), *Psd*/*Aco2* (orange, 33/107, 30%), *Psd*/*Pum2* (cyan, 5/107, 4.6%) and *Psd*/FMRP (white, 45/107, 42%). The third column shows the intersect of *Psd* colocalized at random (rotated 90 degrees) (*Psd*/*Camk2a* = 56/107, 52%; *Psd*/*Aco2* = 25/107, 23%; *Psd*/*Pum2* = 3/107, 2.8%; *Psd*/FMRP = 32/107, 29%). Scale = 5 µm. **C.** Correlation plot of percent *Psd* colocalized with the other 11 mRNAs after subtraction of random colocalization and their abundance (R^2^ = 0.92). **D.** Pie chart of *Psd* mRNA puncta compositions. The data shown here are averaged from four 52 X 52 µm^2^ ROIs per animal then averaged across N=4 mice. 91.86 ± 1.8% (vs. 65.1 ± 5.4% random in Supp. Fig. 7) of *Psd* RNA have overlapping puncta from all other mRNA channels combined, i.e., colocalized mRNAs that include dimers (only one other mRNA) and multimers (at least two other mRNAs). The percentage of colocalized *Psd*-dimers is almost at a level that was observed at random for each pair, except for *Camk2a*, where the colocalization is higher than random (11.58 ± 3.1% of *Psd-Camk2a* dimers vs. random 6.68 ± 1.1%). 57 ± 5.3% of *Psd* mRNA puncta are multimers that have *Camk2a* and at least one other colocalized mRNA, which is higher than random (34.89 ± 4.9%). 12.34 ± 2.9% of *Psd* mRNA are also multimers (vs. random 8.29 ± 1.6%) but do not have *Camk2a*. Lastly, 8.13 ± 1.8% of *Psd* mRNA are not colocalized with any of the other mRNAs in our dataset, which is noticeably lower than observed at random (34.9 ± 5.4%). (N=4 mice.)

Next, we hierarchically clustered the shared colocalization patterns, which revealed three distinct clusters displaying consistently “high”, “intermediate”, or “low” levels of colocalization across all pairwise comparisons. Unexpectedly, levels of colocalization increased with mRNA abundance such that the three most abundant mRNAs in our dataset, *Camk2a*, *Ddn* and *Dlg4* (# of RNA puncta: *Camk2a* = 12,829 ± 1,646; *Ddn* = 11,114 ± 1,262, *Dlg4* = 6,426 ± 424, N=4 mice, Supp. Fig. 4) exhibited uniformly high colocalization patterns with each of the other mRNAs. mRNAs with intermediate levels of abundance (*Calm1* = 6,451 ± 2,096, *Aco2* = 5,054 ± 450, *Psd* = 3,648 ± 764; N=4 mice, Supp. Fig. 4) consistently demonstrated intermediary levels of colocalization across all pairwise comparisons. mRNAs with relatively lower levels of abundance (*Pld3* = 3,457 ± 427, *Cyfip2* = 3,919 ± 725, *Adcy1* = 2,327 ± 407, *Bsn* = 3,157 ± 450, *Pum2* = 1,694 ± 208, *Ppfia3* = 1,162 ± 178, N=4 mice, Supp. Fig. 4) typically showed lower levels of colocalization across all pairwise comparisons.

When pairwise colocalization was analyzed as a percentage of both mRNAs in the pair, the hierarchical clustering similarly scaled by mRNA abundance (Supp. Fig. 6). The degree of colocalization ranged from 2.8% (*Adcy1/Ppfia3*) to 24.5% (*Camk2a/Ddn*). In both analyses, the most abundant mRNAs (*Camk2a*, *Ddn*, *Dlg4*) colocalized the most and the least abundant mRNAs (*Ppfia3*, *Pum2*) colocalized the least across all pairwise comparisons. The intermediary expressors were more variable in their specific order, but followed a similar trend. Representative images of high (*Camk2a*), intermediate (*Aco2*) and low (*Pum2*) levels of colocalization with *Psd* mRNA are shown in Fig. 3B, including the intersecting pixel overlaps for properly registered “experimental” and rotated “random” images. We chose *Psd* as an example mRNA due to its intermediate level of abundance, high % colocalization with *Camk2a*, and its broad fluorescent puncta area distribution presumably reflective of multiple populations of homotypic RNA particles. To visualize the influence of mRNA abundance on colocalization with the other 11 mRNAs, we plotted the percent *Psd* colocalization above random versus abundance in a correlation plot (Fig. 3C). We found that the abundance of the mRNAs is highly correlated with the percent *Psd* colocalization (R^2^= 0.92). We observed the same pattern regardless of the expression of the mRNA that is being colocalized to, as *Ddn* (high expressor) and *Pum2* (low expressor) colocalization values were also highly correlated with abundance (*Ddn* R^2^= 0.98*, Pum2* R^2^= 0.95, Supp. Fig. 7).

Considering the technical limitation that pairwise colocalization values cannot portray the colocalization of multiple (more than two) RNAs, we then quantified the percentage of *Psd* RNA that are localized in association with at least one (dimers) or more RNA species (multimers) in properly registered experimental and rotated random images (Fig. 3D and Supp. Fig. 8). Because the sum of measured pairwise colocalization of *Psd* mRNA (Supp. Fig 5A) exceeded 100%, it implied that a percentage of *Psd* mRNA was multimers (more than two mRNAs colocalized). The pie chart of *Psd* mRNA compositions from experimental images shows that 8.13 ± 1.8% of *Psd* mRNAs are not colocalized with any mRNA in our dataset (singleton, Fig. 3D), which is lower than observed for the 90-degree rotated random image (34.9 ± 5.4%, Supp. Fig. 8). This suggests that a large fraction of the *Psd* mRNAs are heterotypic mRNA puncta, containing different types of mRNA transcripts, including *Psd*-dimers, which have *Psd* colocalized with only one other type of transcript (measured here) and *Psd*-multimers which contain more than two different transcripts including *Psd* (Fig. 3D and Supp. Fig. 8). The percentage of *Psd*- dimers range from 0.23 ± 0.1% (*Psd*/*Ppfia3*) to 11.58 ± 3.1% (*Psd*/*Camk2a*) which mirrored the abundance of the mRNAs. However, only the percentage of *Psd/Camk2a* dimers (11.58 ± 3.1%) is higher than random (6.68 ± 1.1%); all other *Psd*-dimers are near random levels. We observed that 70.25 ± 4.8% of *Psd* RNAs have at least three or more (including *Psd*) transcripts (multimers) of which the greatest fraction has *Camk2a* (57.91 ± 5.3%). Only 12.34 ± 2.9% of multimer *Psd* RNAs were without *Camk2a*, underscoring the dominating presence of *Camk2a* in both *Psd* dimers and multimers in our data. In contrast, the percentage of *Psd* multimers with (34.89 ± 4.9%) and without *Camk2a* (8.29 ± 1.6%) are lower in the rotated random images. Furthermore, when other mRNAs were quantified similarly, we found that an average of 88.6 ± 0.73% of each neuropil localized mRNA is localized with at least one other mRNA in our dataset compared to 65.62 ± 0.66% observed at random (all 12 comparisons are significantly higher than random, unpaired two sample Welch’s t-test with FDR correction) (Supp. Fig. 9). Only ∼11.40 ± 0.73% of the mRNAs are not colocalized with any of the other 11 mRNAs in this dataset. To confirm that the anatomical orientation of the neuropil has no effect on the calculation of random colocalization, we also rotated each mRNA image 180 degrees to calculate the percentage of random overlapping puncta and did not observe any noticeable difference from 90 degrees (Supp. Fig. 9). However, we note the important caveat that there are assumed anatomical constraints (areas unavailable for colocalization) that are missing in the rotated image comparison that make these colocalization values an underestimate.

To test whether association with FMRP influences the relationship of pairwise colocalization or heterotypic RNP associations, we subsetted the dataset by only selecting the mRNA puncta from each channel that colocalized with FMRP protein (Supp. Fig. 10). Our hypothesis was that the observed colocalization after random subtraction would be higher in the presence of FMRP indicating that having a shared multivalent transacting protein makes mRNAs more likely to multimerize with each other into common RNPs. In properly registered experimental images, the degree of pairwise mRNA-mRNA colocalization in FMRP containing RNPs ranged from 2.8% (*Pld3/Ppfia3*) to 60.3% (*Psd/Camk2a*) (Supp. Fig. 10Ai). We then, as mentioned above, calculated pairwise colocalization by rotating one image from each mRNA pair to 90 degrees (Supp. Fig. 10Aii). Random pairwise colocalization ranged from 0.3% (*Adcy1/Ppfia3*) to 16.6% (*Adcy1/Camk2a*). After random colocalization was subtracted from the experimental colocalization, the percentage of pairwise colocalization ranged from 1.8% (*Pld3/Ppfia3*) to 47.4% (*Psd/Camk2a*), revealing a similar trend of pairwise colocalization that tracks with mRNA abundance (Supp. Fig. 10B). Although the general trend was similar, we saw an overall gain of % colocalization for each pair in the FMRP-containing RNPs compared to the computation including all RNAs as shown by the correlation plot comparing *Psd*-pairwise % colocalization with and without FMRP (Supp. Fig. 10C).

We then quantified the percentage of FMRP-containing *Psd* mRNAs that are colocalized with at least one (dimer FMRP cargo) or more species of mRNAs (multimer FMRP cargo) from both properly registered experimental and 90 degree rotated random images. These data are plotted as experimental and random *Psd* RNP composition pie charts (Supp. Fig. 10DE). As we noted previously for all mRNAs detected, the majority of FMRP-containing *Psd* mRNAs (86.4%) are colocalized with at least one other mRNA along with FMRP protein, which is higher than random (25.3%). The difference in experimental versus random colocalization (∼3.3 fold) is greater than reported for analyses without FMRP (∼1.3-fold, Supp. Fig 9), driven by a decrease in the random but not experimental values. Of these *Psd*-FMRP cargoes, the percentage of *Psd* having only one other mRNA (*Psd*-dimers) is at the level of random colocalization except for *Psd*-*Camk2a* (22.6% vs random 5.1%). 41.06% *Psd*-FMRP cargoes are multimers with *Camk2a* mRNA as opposed to only 7.8% observed at random, indicating the predominance of *Camk2a* in both *Psd*-FMRP-dimers and *Psd*-FMRP-multimers. 8.8% of *Psd*-FMRP cargoes are also multimers, higher than random (3.8%), that have three or more mRNAs without *Camk2a*. Lastly, 13.6% of FMRP-containing *Psd* RNPs are segregated from the other 11 mRNAs, which is remarkably lower than observed at random (74.6%). These data suggest that for these 12 neuropil localized mRNAs, when associated with FMRP, have a higher likelihood than random to multimerize with each other, with a bias for the highly abundant ones.

### Variation in neuropil RNA abundance is sufficient to scale mRNA colocalization across cell types

To further explore the relationship of mRNA colocalization with mRNA abundance, we examined whether differences in the abundance of one mRNA within a pair (across different cell types) influenced the colocalization patterns of that mRNA. We performed smFISH for *Rgs14*, *Adcy1* and *Ppp1r9b* which are known hippocampal dendritic mRNAs (Farris et al., 2019) downstream of group I metabotropic glutamate receptor (mGluR1/5) Gq-mediated signaling (Evans et al., 2018; Morris et al., 2023; Samadi et al., 2023; H. Wang et al., 2008; H. Wang & Zhuo, 2012; X. Wang et al., 2007). Consistent with previous observations, all three mRNAs localized throughout the neuropil, both proximal and distal layers, in CA2 and CA1 cell types as well as the molecular layer of dentate gyrus (DG) (Fig. 4AB). However, each mRNA has a distinct expression pattern reflected by differences in the number of mRNA puncta, which is consistent with previous RNAseq studies (Farris et al., 2019; Hale et al., 2021). mRNA puncta number varies from hundreds (*Rgs14*) to thousands (*Adcy1* and *Ppp1r9b*) of transcripts (Fig. 4C). When comparing the same sized region of neuropil, the number of *Ppp1r9b* mRNA was similar in CA2, CA1 and DG (# of *Ppp1r9b* DG: 8579 ± 1685, CA1: 6679 ± 1619, CA2: 9409 ± 1881, no effect of cell type, oneway repeated measures analysis of variance [RM ANOVA]: F = 4.566, p = 0.132, N=4 mice). *Rgs14* mRNA count was significantly higher in CA2 (# of *Rgs14* mRNA DG: 204 ± 30, CA1: 196 ± 28, CA2: 490 ± 145, significant effect of cell type, RM ANOVA: F = 21.36, p = 0.0030) than CA1 (p = 0.0306, Tukey’s post hoc test) and DG (p = 0.0191, Tukey’s post hoc test). *Adcy1* mRNA count was almost 2.5-fold higher in DG (# of *Adcy1* DG: 2969 ± 501, CA1: 943 ± 187, CA2: 1202 ± 180, significant effect of cell-type, RM ANOVA: F = 108.3, p = 0.0007) than CA1 (p = 0.007, Tukey’s post hoc test) and CA2 (p = 0.0018, Tukey’s post hoc test).

**Fig 4.**
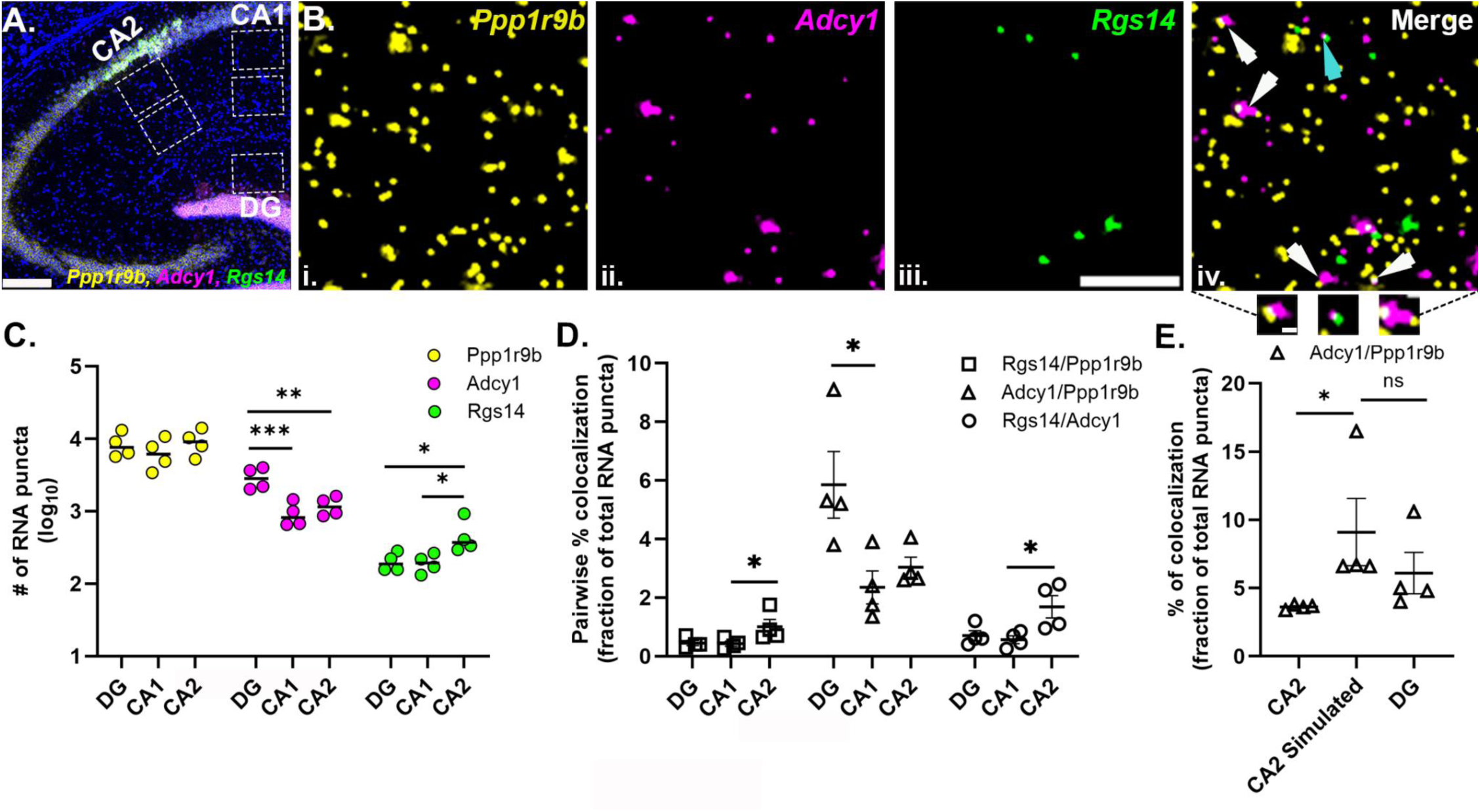
Neuropil mRNA colocalization patterns scale with mRNA abundance across cell types with varied mRNA expression. **A.** Representative tilescan image of *Ppp1r9b* (yellow), *Adcy1* (magenta) and *Rgs14* (green) mRNAs and nuclei (blue) in adult mouse hippocampus. Dashed white boxes are regions analyzed from CA1 and CA2 proximal and distal dendrites and DG. **B.** High-magnification representative images of (i) *Ppp1r9b*, (ii) *Adcy1*, (iii) *Rgs14* and (iv) merged in CA2 distal dendrites. Arrows indicate colocalization of *Adcy1* with either *Ppp1r9b* (white) or *Rgs14* (cyan). Three example images of RNA colocalization in the callout section. **C.** Quantification of the number of *Ppp1r9b*, *Adcy1* and *Rgs14* RNAs in DG, CA1 and CA2. (# of *Ppp1r9b* mRNA in DG: 8579 ± 1685, CA1: 6679 ± 1619, CA2: 9409 ± 1881, one-way repeated measures analysis of variance [RM ANOVA]: F = 4.566, p = 0.132) (# of *Adcy1* mRNA in DG: 2969 ± 501, CA1: 943 ± 187, CA2: 1202 ± 180, one-way repeated measures analysis of variance [RM ANOVA]: F = 108.3, p = 0.0007) (# of *Rgs14* mRNA in DG: 204 ± 30, CA1: 196 ± 28, CA2: 490 ± 145, one-way repeated measures analysis of variance [RM ANOVA]: F = 21.36, p = 0.0030). Stats were run on the transformed (log_10_) values as plotted to meet the normality assumption. Tukey’s post hoc tests reported on the plot. **D.** Pairwise % colocalization of *Rgs14*/*Ppp1r9b*, *Adcy1*/*Ppp1r9b* and *Adcy1*/ *Rgs14* expressed as a percentage fraction of total number of RNAs in the pair. Some comparisons failed the Shapiro-Wilk normality test so a Friedman rank based ANOVA was run. *Adcy1/Ppp1r9b* Friedman statistic = 8.00, P = 0.0046, *Adcy1/Rgs14* Friedman statistic = 6.50, P = 0.041, *Rgs14/Ppp1r9b* Friedman statistic = 6.50, P = 0.041, Dunn’s post hoc test reported on the plot. **E.** *Adcy1*/*Ppp1r9b* pairwise colocalization in experimental CA2 and DG images compared to simulated CA2 images where *Adcy1* RNA puncta were randomly added to make CA2 equivalent to DG. Data failed the Shapiro-Wilk normality test so a Friedman rank based ANOVA was run. Friedman statistic = 8.00, P = 0.0046. Dunn’s post hoc test reported on the plot. N=4 mice. Error bars indicate SEM. *P<0.05, **P<0.01, ***P<0.001; Scale bars: A) 200µm, B) 5µm, 0.5µm.

We then tested whether differences in mRNA expression influences mRNA pairwise colocalization values across these three cell types (Fig. 4D). We quantified the number of overlapping puncta (>1% overlap) between two channels and expressed that number as a percentage of the total number of mRNA puncta in the pair (Fig. 4D). Based on our HiPlex data, we predicted that the *Adcy1*/ *Ppp1r9b* mRNA pair would demonstrate higher colocalization compared to other pairs (*Adcy1/Rgs14*, *Rgs14/Ppp1r9b*) in all three cell types and this effect would be significantly higher in DG compared to that in CA2 and CA1 due to the significantly higher number of *Adcy1* mRNA in DG. Indeed, we observed 5.85 ± 1.14% of total *Adcy1*/*Ppp1r9b* mRNA were colocalized in DG compared to 2.35 ± 0.56% in CA1 and 3.04 ± 0.34% in CA2 (Significant effect of cell type, rank based ANOVA, Friedman statistic: 8.00, p=0.0046, N=4 mice) and this difference was significant for DG vs CA1 (p=0.0140, Dunn’s multiple comparison test). Similarly, we also predicted that *Adcy1/Rgs14* would be highly colocalized in DG compared to CA1 and CA2 due to the higher abundance of *Adcy1* in DG (Significant effect of cell type, rank-based ANOVA, Friedman statistic: 6.500, p=0.0417). Instead, *Adcy1*/*Rgs14* demonstrated higher colocalization in CA2 (1.69 ± 0.38%), which has the highest number of *Rgs14* mRNA among these three cell types, and this effect was significant compared to CA1 (colocalized *Adcy1/Rgs14* = 0.56 ± 0.13%, p=0.0400, Dunn’s post hoc test, N=4 mice). Restricting colocalization to >50% overlap (versus 1%) in between two channels did not change the relationship of mRNA colocalization with abundance in our dataset (Supp. Fig 11). These data indicate that mRNA abundance similarly influenced colocalization in each of the three cell types we analyzed.

We then tested whether simulating an increase in mRNA abundance would reproduce the expected increase in pairwise colocalization. Specifically, we randomly added *Adcy1* mRNA puncta to CA2 images to make them have the equivalent number of *Adcy1* puncta as DG images taken from the same brain section, while keeping the number of other two mRNAs constant (# of *Adcy1* in DG: 2973 ± 726, CA2 proximal: 1248 ± 106, CA2 simulated: 2973 ± 726; # of *Ppp1r9b* in DG: 8346 ± 1449, CA2 proximal: 8840 ± 1698, CA2 simulated: 8840 ± 1698; # of *Rgs14* in DG: 235 ± 37, CA2 proximal: 622 ± 175, CA2 simulated: 622 ± 175, N=4 mice). The increased *Adcy1* mRNA in simulated CA2 resulted in a significant increase in pairwise % colocalization of *Adcy1*/*Ppp1r9b* mRNAs similar to experimental values in DG (% *Adcy1*/*Ppp1r9b* colocalized in CA2: 3.62 ± 0.09%, DG: 6.1 ± 1.51%, Simulated CA2: 9.1 ± 2.47%; significant effect of cell type, Friedman statistic = 8.00, P = 0.0046, N= 4 mice; Dunn’s post hoc tests: CA2 vs CA2 simulated: p = 0.0140, DG vs CA2 simulated: p = 0.4719). The slightly higher *Adcy1/Ppp1r9b* % colocalization in simulated CA2 compared to DG is likely due to the increase in *Ppp1r9b* mRNA puncta count in the chosen ROIs compared to DG. Thus, the most likely explanation of the observed colocalization is due to random overlaps driven by spatial proximity of mRNAs which in turn is dictated by mRNA abundance.

Lastly, building on our previous observation of fluorescent mRNA puncta area heterogeneity, we also quantified the fluorescent mRNA puncta area distributions of these mRNAs in CA1 and CA2 proximal and distal neuropil layers and DG. We found no differences between proximal and distal layers for puncta area and therefore the data were collapsed and represented as one neuropil population per animal for CA2 and CA1. Across animals, we consistently observed the same pattern of puncta area heterogeneity for each mRNA species in all three hippocampal cell types (Supp. Fig. 12). However, the distributions of mRNA puncta area were quite heterogeneous across mRNAs; in particular, *Rgs14* mRNA puncta were consistently small sized puncta in all three cell types with their largest relative percent peak at 0.1 µm^2^ (68.5 ± 3.1%, averaged across mice for each cell type and then averaged across cell types). *Adcy1* and *Ppp1r9b*, however, showed a large broad distribution with the largest relative percent peak at 0.20 um^2^ (*Adcy1:* 34.2 ± 3.1%; *Ppp1r9b*: 27.9 ± 1.1 %) and a qualitatively distinct larger population of mRNAs that are greater than 0.5 um^2^ (*Adcy1*: 5.35 ± 2.04%; *Ppp1r9b*: 10.7 ± 1.2%). These large mRNA particles were not present for *Rgs14* mRNA (*Rgs14* median puncta area CA1: 0.10 ± 0.01 µm^2^, CA2: 0.10 ± 0.02 um^2^, DG: 0.10 ± 0.02 um^2^; no effect of cell-type, RM ANOVA, F = 0.026, p = 0.9744, N=4 mice). *Adcy1* mRNA median puncta area CA1: 0.22 ± 0.03 µm^2^, CA2: 0.22 ± 0.03 µm^2^, DG: 0.19 ± 0.02 µm^2^; no effect of cell type, RM ANOVA: F = 0.5406, p = 0.6083, N=4 mice; *Ppp1r9b* median puncta area CA1: 0.22 ± 0.01 µm^2^, CA2: 0.24 ± 0.02 µm^2^, DG: 0.22 ± 0.03 µm^2^; no-effect of cell type, RM ANOVA, F = 0.5507, p = 0.6032, N=4 mice). Compared to the area distribution in our HiPlex data, the histogram of mRNA puncta area in this experiment was skewed smaller although the relative distinction between “small” and “large broad” clusters based on fluorescent puncta area were consistent. We compared the median puncta area of *Adcy1* mRNA in the CA2 distal neuropil from this experiment with the HiPlex CA2 distal neuropil images and found no significant difference between Adcy1 median puncta area between the two experiments (*Adcy1* median puncta area in CA2 distal neuropil measured in HiPlex smFISH: 0.25 ± 0.01, and 3Plex smFISH: 0.24 ± 0.04, two-tailed unpaired t-test, p = 0.7304). These data suggest that these mRNAs can exist in consistently similar sized populations (both small and large homotypic RNA complexes) across multiple cell types despite endogenous differences in their expression.

## DISCUSSION

In this study, we visualized 15 neuropil localized mRNAs to investigate how mRNAs are spatially organized for delivery to synapses in intact rodent hippocampus. First, we provide evidence supporting the heterogeneity of neuropil mRNAs by describing differences in mRNA fluorescent puncta area. We interpret this data to reflect differences in the amount of individual mRNA transcripts per mRNA smFISH puncta. Second, by simultaneously visualizing a dozen neuropil localized FMRP target mRNAs, we found that every mRNA we investigated, regardless of its abundance, colocalizes more with the highly abundant mRNAs compared to the lower abundance mRNAs. This result stands after correcting for random colocalization or the total fraction of the two mRNAs being compared. Our findings were similar for RNPs defined by the presence of FMRP. Third, the data suggest that mRNA colocalization correlates with mRNA abundance across multiple hippocampal cell types, an effect that can be recapitulated by simulations of mRNA abundance. Thus, we failed to identify selectivity in how these mRNAs associate with each other in the neuropil. Instead, the probability of these mRNAs interacting within the neuropil appears to be stochastic and linked to their neuropil abundance.

### Localized mRNAs contain varying amounts of a single mRNA species

Neurons localize thousands of different mRNAs of variable abundance and subcellular distributions to support synaptic function. Yet, few studies have systematically characterized how mRNAs in the hippocampal neuropil are sorted into RNPs and how their compositions *in vivo* could support the delivery of thousands of mRNAs encoding proteins involved in many different biological processes. Mikl and colleagues (Mikl et al., 2011) investigated the localization of *Map2* and *Camk2a* mRNAs in hippocampal neurons in culture, showing that these mRNAs are present in dendrites in distinct RNPs, each containing as few as one or only a few copies of the same transcript with minimal colocalization between the two transcripts. Another study by Batish and colleagues (Batish et al., 2012), visualized pairwise combinations of eight dendritically localized transcripts with smFISH in hippocampal cultured neurons, also showing unimodal distribution of mRNA puncta fluorescence intensities and ∼4% of colocalization between pairs of mRNAs, suggesting that mRNA molecules are trafficked singly and independently of others in neurons. In addition, there is evidence from in situ studies assessing individual mRNA content that supports the idea that mRNAs localize in variable copy number states. Single molecule FISH detected β-actin mRNA in live hippocampal cultured neurons showed that RNPs may contain single as well as multiple copies of β-actin mRNA and the copy number decreased with increasing distance from the cell soma (Park et al., 2014). Variations in size and intensity of individual *Camk2a*, *Arc* and neurogranin (*Ng)* RNA granules in developing neurons (fixed) were also reported by Gao et al. (Gao et al., 2008). Further, a recent study by Donlin-Asp and colleagues (Donlin-Asp et al., 2021) used molecular beacons in cultured neurons to individually track endogenous mRNAs by live cell imaging, *Camk2a* and *Psd95*. In addition to detecting single mRNA transport events, they observed mRNA-mRNA fusion events within the same transcript resulting in heterogeneous copy number states in neuronal dendrites. While extremely informative, most of these studies were done in primary neuronal cultures and limited in the number of species of localized mRNAs investigated, demonstrating a need to evaluate RNA copy number and composition for the growing list of localized mRNAs in intact neuronal circuits.

Our data on neuropil localized mRNA puncta sizes (measured as mRNA fluorescent puncta area) in fixed rat and mouse hippocampus (DG, CA1, CA2), corroborate previous observations in culture that mRNA content varies from low-copy number mRNAs to higher order homotypic mRNA clusters of the same transcript (multiple copies of the mRNA derived from the same gene). For the Arc dilution experiment, the decrease in Arc puncta intensity, diameter, and count suggest there are multiple pools of Arc puncta with different amounts of Arc mRNA. However, these data are relative to saturating probe conditions (1X), not to puncta with a known RNA copy number, so we cannot conclusively state that the loss of Arc puncta equates to a loss of puncta containing a single RNA. The differences in puncta size distributions across RNAs was an unexpected finding that became obvious when visualizing a dozen RNAs simultaneously in the same tissue section. However, without similar dilution calibration curves for each RNA and/or fiduciary standards to control for different fluorophores and image acquisition parameters across RNAs, we are not able to make strong claims about the significance of the observed differences. We did not observe any particular round of imaging or fluorophore wavelength to behave in a certain way that would explain the observed differences in puncta area distributions though. We also cannot exclude the possibility that the differences in fluorescent mRNA puncta intensity, diameter, or area could be due to differences in the number of probes bound to a target RNA. However, because we compare puncta size distributions, which include all possible probe intensities/sizes, it is unlikely for different transcript probes (which all have 20 ZZ probe pairs) to result in different distributions unless there is something about the specific transcript (secondary structure etc.) driving the bias. Further, it is difficult to imagine a technical explanation for observed puncta area differences within the same transcript, unless there is a biological mechanism restricting probe access to specific populations of transcripts that results in a non-gaussian distribution of sizes.

Our evidence in support of heterogeneous copy number mRNAs within and across 15 neuropil localized mRNAs is consistent with the idea that multiple RNA assembly states coexist for localizing at least these mRNAs, which may provide flexibility in regulating synaptic activity-induced changes in translation (Fernandez-Moya et al., 2014). Observations from non-neuronal systems, such as drosophila mRNA germ granules (Niepielko et al., 2018; Trcek et al., 2020) also indicate that localized mRNAs sort into homotypic clusters. However, with the limited number of visualized neuropil localized mRNAs, it is not yet clear whether the existence of distinct copy number states is a transcript-specific feature or a transcriptome-wide phenomenon. mRNA constituents of transporting or localized mRNA granules identified by synaptoneurosome- or brain lysate-fractionation are present in monosomes as well as in translationally silent stalled polysomes (Hafner et al., 2019; Kanai et al., 2004; Krichevsky & Kosik, 2001). Therefore, whether different sized mRNAs in our dataset reflect functional differences such as their association with other mRNAs and/ or ribosomes or translational status is yet to be determined. Work on other types of cytoplasmic granules (p-bodies and stress granules) in living cell lines, show that granule size correlates with increased granule stability (Moon et al., 2019). Further studies are needed to identify whether differences in mRNA size and composition reflect different structural properties and/or functional mRNA states.

### mRNA colocalization within the neuropil scales with mRNA abundance

The composition of certain types of specialized mRNA granules (e.g. stress granules, germ granules) is influenced by mRNA abundance (Bauer et al., 2022; Trcek et al., 2020; Van Treeck et al., 2018). Data on localized neuronal RNAs, however, in the context of being influenced by mRNA abundance is comparatively limited. Biophysical studies provide evidence that mRNA concentration, in addition to mRNA structure and stability, favors in vitro mRNA-protein condensate formation (Boeynaems et al., 2019; Garcia-Jove Navarro et al., 2019; Roden & Gladfelter, 2021). Experiments in *Drosophila* germ cell granules show that highly abundant mRNAs have higher seeding events to initiate homotypic RNP formations through self-recruitment and subsequently recruit other mRNAs to the RNPs (Niepielko et al., 2018). Our pairwise colocalization data, using p17 mouse brain, shows that highly abundant neuropil mRNAs (*Camk2a, Ddn, Dlg4*) are spatially distributed such that they colocalize the most with the other neuropil mRNAs in our dataset, reflecting the higher availability of these transcripts to possibly promote clustering with other mRNAs into heterotypic RNPs. However, it is important to note that our data rely on a colocalization metric based on 2D overlapping in situ signals limited by 250 nm x-y resolution and a low overlap cut off of 1%, which likely overestimates the true level of colocalization. Though the effect of abundance remained when the overlap was more stringent at 50%. Nevertheless, additional super-resolution techniques (i.e. STORM) are required to prove whether any mRNAs investigated in this study are physically clustering (<250 nm).

Analysis of the data as a percentage of total mRNA puncta in the pair showed that 24.5 % of *Camk2a* and *Ddn* (two highest abundant in our dataset) are colocalized in the neuropil whereas, only 3.5% of *Pum2* and *Ppfia3* (two least abundant in our dataset) are colocalized in the neuropil. Consistently, Wang & colleagues visualized 950 mRNA transcripts in neuronal dendrites in culture (18 days in vitro) using the barcode-based imaging method, MERFISH (Multiplexed Error-Robust Fluorescence In Situ Hybridization), combined with expansion microscopy and spatial proximity clustering to show that dendritic mRNAs with high abundance spatially cluster together (*Camk2a, Ddn, Dlg4, Ppp1r9b, Shank1, Palm*) (G. Wang et al., 2020). It is plausible to hypothesize that highly abundant dendritic mRNAs may have a greater chance of being present in numerous small RNPs that then assemble into larger mRNA granules. However, we cannot rule out whether other mRNA-specific features, in addition to their high abundance, may also be contributing to the observed high levels of colocalization. For example, *Camk2a, Ddn*, *Dlg4* and *Ppp1r9b* transcripts, in addition to being highly abundant in the neuropil, are also the mRNAs that are highly translated in the dendrite as shown by their high ribosomal densities (Glock et al., 2020). *Camk2a* and *Dlg4* have been predicted by in silico tools to be strongly interacting with FMRP (Cirillo et al., 2013), possibly indicating that certain features specific to these mRNAs, other than their high expression, may also contribute to their noticeable presence in the majority of neuropil localized mRNA clusters in our data. Moreover, studies in neurons and non-neuronal cell types have shown that mRNA localization is influenced by several sequence-specific features including, but not limited to, sequence length, RBP-binding motifs, and other cis sequences influencing stability (Farris et al., 2019; Lipshitz & Smibert, 2000; Middleton et al., 2019; H. Wu et al., 2020). Thus, further investigation into the sequence-specific features of the 15 neuropil localized mRNAs in our dataset is needed to determine whether and/or how highly abundant FMRP target and non-target mRNAs cluster with other classes of mRNAs in the neuropil.

### Stochastic mRNA interactions based on spatial distributions

Observations in previous smFISH studies have reported conflicting conclusions on mRNA colocalization in dendrites (Batish et al., 2012; Gao et al., 2008; Mikl et al., 2011; Tübing et al., 2010). The percentage of colocalized mRNA molecules in pairwise combinations ranged from 0.33 - 3.36% (Batish et al., 2012), 5.7 - 8.3% (Mikl et al., 2011), 7 - 32% (Farris et al., 2014) depending on the quantitative definition of colocalization, analysis technique, as well as the mRNA combinations that were compared. However, it is unclear 1) whether these observations took into account the colocalization of transcripts that would occur randomly and 2) whether these interpretations (i.e., predominantly single mRNA transport) would remain valid if more than a few molecules were visualized at once. Our HiPlex and 3-color smFISH experimental images as well as simulated data shows pairwise colocalization of mRNAs, calculated as percentage of each mRNA, ranging from ∼1% to ∼30% after subtraction of random colocalization. Since pairwise colocalization is not sufficient to map the extent of multi-transcript RNA granule composition, we then quantified the percentage of a given mRNA that localizes with any of the other eleven mRNAs. We found that ∼90% (∼65% random) of any mRNA species that we examined are colocalized with at least one other RNA, which would suggest mRNA co-distribution is the favored form of localization for these mRNAs. However, we need to be careful in our interpretation of this data, as image rotation does not fully recapitulate the cytoarchitecture (e.g., extracellular spaces, organelles, etc.) present in properly registered images that theoretically restricts the potential locations for overlap, which will result in an underestimation of random colocalization, thereby inflating the random-subtracted colocalization percentage observed. This is evident in the pairwise comparisons near random that are always above 0. To circumvent this, we performed analyses to simulate true random colocalization of specific RNA pairs, which resulted in higher colocalization values, although this might not relate to other gene combinations that have different spatial distribution patterns. Further, when we subsetted instances of colocalization to only those defined by the presence of FMRP protein, the percentage of multi-RNA containing clusters (86.4%) were similar to the percentages without FMRP (91.9%), however, the percentages detected at random were considerably decreased (25.3%), indicating the robustness of assessing multi-RNA containing clusters compared to pairs.

Whether RNAs are independently segregated or co-distributed into common complexes has been debated in the field for many years. Interestingly, as was proposed in the neuronal mRNA-transport sushi-belt model, mRNAs patrol the neuronal processes in a multidirectional fashion with intermittent rest and run times and dynamic transient interactions (Ahn et al., 2023; Bauer et al., 2019; Song et al., 2018). Such a model would then predict, highly abundant mRNAs have a higher likelihood of random transient interactions during localization in a cell-autonomous fashion and the precision of sorting is obtained locally at the synapse level. As synaptic activity has been shown (experimental and modelling studies) to influence the random oscillatory behavior of mRNAs in transport, further characterization is critical to understand how our snapshot observation of mRNA spatial distribution driven stochastic mRNA colocalization changes in response to local cues (Buxbaum et al., 2015). There are multiple lines of evidence that show FMRP targets are differentially altered (or not) in the absence of FMRP at the level of mRNA localization (Dictenberg et al., 2008; Miyashiro et al., 2003; Steward et al., 1998). Future perturbation experiments are required to assess whether FMRP-containing RNP compositions make the mRNA cargoes more or less vulnerable to FMRP loss. Moreover, there is compelling evidence of FMRP granule remodeling after synaptic activity to support local protein synthesis (Kharod et al., 2023; Sidorov et al., 2013) (Kharod et al., 2023; Sidorov et al., 2013). Further investigation is needed to determine how selectivity and specificity is achieved for FMRP-containing RNPs different types of mRNA transcripts in order to appropriately remodel the synaptic proteome.

In particular, *Camk2a* mRNA has been shown to interact with multiple other RBPs in addition to FMRP (RNG105 (Nakayama et al., n.d.; Shiina et al., 2005), CPEB (L. Wu et al., 1998), and Staufen (Ortiz et al., 2017). In our data, *Camk2a* was dominating the spatial overlaps with other mRNAs whether they contained FMRP or not. It seems reasonable to hypothesize that *Camk2a*-containing heterotypic RNPs might achieve some degree of mRNA selectivity based on co-regulation with *Camk2a*-associated RBP(s). We were unable to investigate this relationship with the limited number of neuropil localized RBP and mRNAs we co-visualized. Indeed, it is certainly possible that our findings of stochastic neuropil mRNA interactions and the relationship with mRNA abundance may not translate to RNPs composed of other mRNAs, or other FMRP target RNAs, or RNPs defined by specific sets of RBPs that may confer specificity not detectable to the methods and analyses used here. By taking advantage of advanced tools in spatial imaging, the results of this study provide descriptive, but not causal, evidence that 15 neuropil localized mRNAs localize in heterogeneous copy number states and exhibit stochastic spatial clustering explained by mRNA abundance-a model predicted by mathematical modelling studies to be energetically cost-efficient (Bergmann et al., 2025; Wagle et al., 2023). Follow up studies are needed to elucidate whether such spatial localization patterns are influenced by RBPs or other factors to functionally influence the localization and translation of messages at synapses.

## Supporting information

HiPlex RNA Probe Information

## Acknowledgements

We would like to acknowledge Dr. Joun Park for contributing to the *Shank2* in situ studies and the investigation into *Adcy1* RNA sizes. We also acknowledge Dr. Gail Lewandowski for contributing to the image acquisition of the Arc probe dilution experiment. The authors acknowledge resources and support from the Virginia Tech animal care staff and Cellular and Molecular Imaging Core, part of the Fralin Biomedical Research Institute at VTC. This research would not be possible without the use of the Nikon Elements software.

## Authors’ contributions

Designed research – SF, OS, SAS; Performed research – RT, GM, FQ, SF; Analyzed data – RT, GM, FQ, SF; Writing Original Draft – RT, SF; Review & Editing – RT, SF, SAS, OS; Funding Acquisition – SF, OS.

---

**Supplemental Fig. 1.**
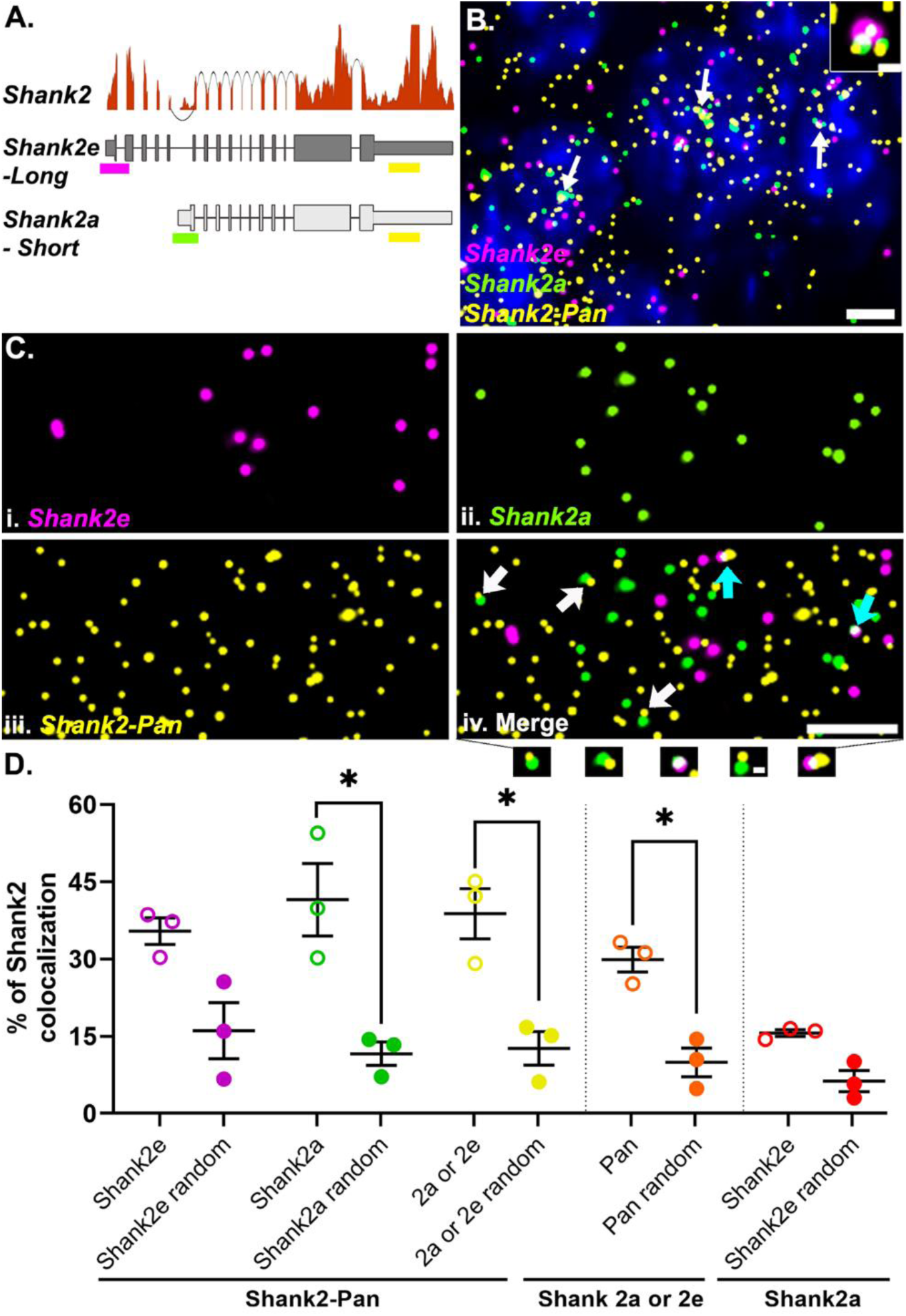
*Shank2* isoform-specific 5’ probes are highly colocalized with the Pan 3’ probe. **A.** *Shank2* isoform gene models with RNAseq read depth data showing the relative expression levels in hippocampal CA2. Sequences from either long (*Shank2e*) or short (*Shank2a*) transcripts targeted by different 5’ probes (magenta and green, respectively) and both targeted by the Pan 3’ probe (yellow) are shown. **B.** Representative image of the three *Shank2* probes in CA2 cell bodies. Nuclei are labeled with DAPI (blue). White arrows indicate example transcriptional foci. Dashed white box is the inset showing a transcriptional focus labeled by all three probes. **C.** High-magnification images of (i) *Shank2e*, (ii) *Shank2a*, (iii) *Shank2*-Pan and (iv) the merged image. Arrows indicate example colocalization of *Shank2*-Pan 3’ probe with either *Shank2e* 5’ probe (cyan arrows) or *Shank2a* 5’ probe (white arrows) as shown below. **D.** Quantification of the % colocalization between the *Shank2e* (magenta) and *Shank2a* (green) or both (yellow) with the *Shank2*-Pan probe (open circles) compared to that observed by random colocalization (closed circles). The percent of *Shank2*-Pan colocalized with either *Shank2a* or *Shank2e* (orange) compared to random colocalization and the percent of *Shank2e* colocalizing with *Shank2a* (red) compared to random, many of which are transcription foci, as shown in B. Error bars indicate SEM; N=3 mice; * denotes p <0.05 from paired one-tailed t-test. Scale bars: B) 5 µm, 1 µm, C) 5 µm, 0.5 µm.

**Supplemental Fig. 2 (Refers to Fig. 2 and 3).**
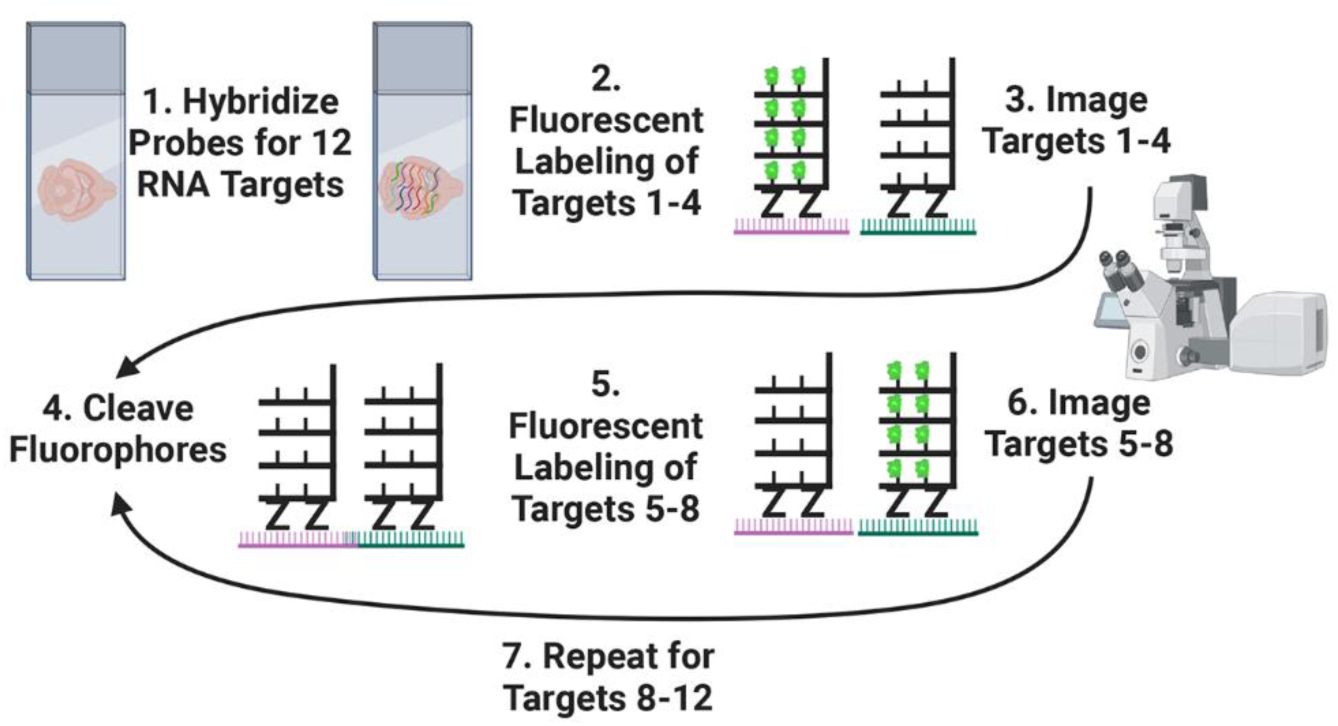
Schematic showing workflow of HiPlex smFISH.

**Supplemental Fig. 3 (Refers to Figs. 2-4).**
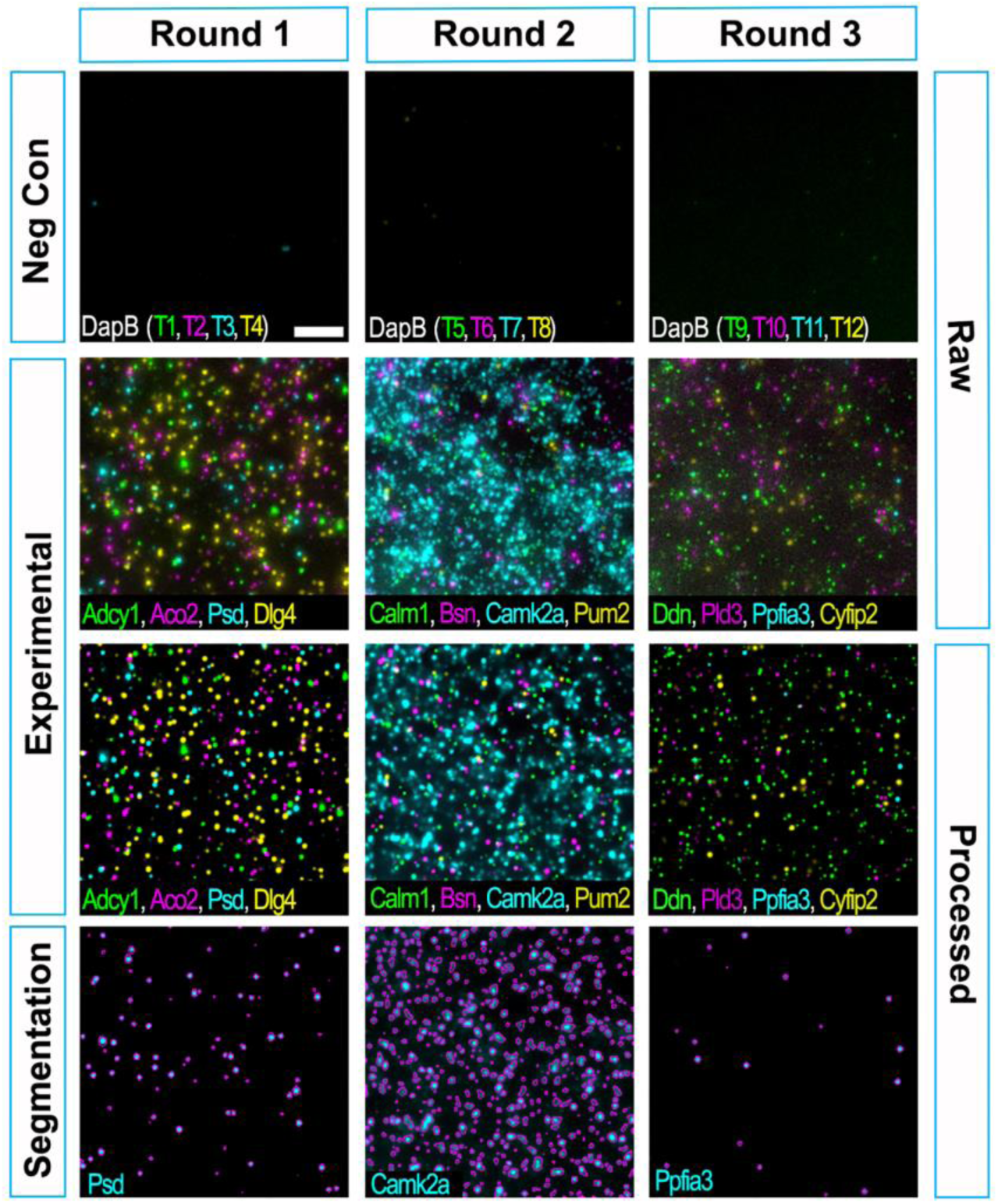
HiPlex image processing and segmentation. Raw and processed negative control images probed for the bacterial RNA *DapB* in each channel. Negative control images were acquired with identical acquisition parameters as experimental images shown below from all three rounds of HiPlex smFISH. Experimental images are presented with the same intensity thresholds as the corresponding negative control channels. The last row displays segmented binary layers for the *Psd, Camk2a*, and *Ppfia3* channels, created using intensity thresholds determined from the negative control image of the corresponding channels in each round. Scale: 5 µm.

**Supplemental Fig. 4 (Refers to Fig. 2 and 3).**
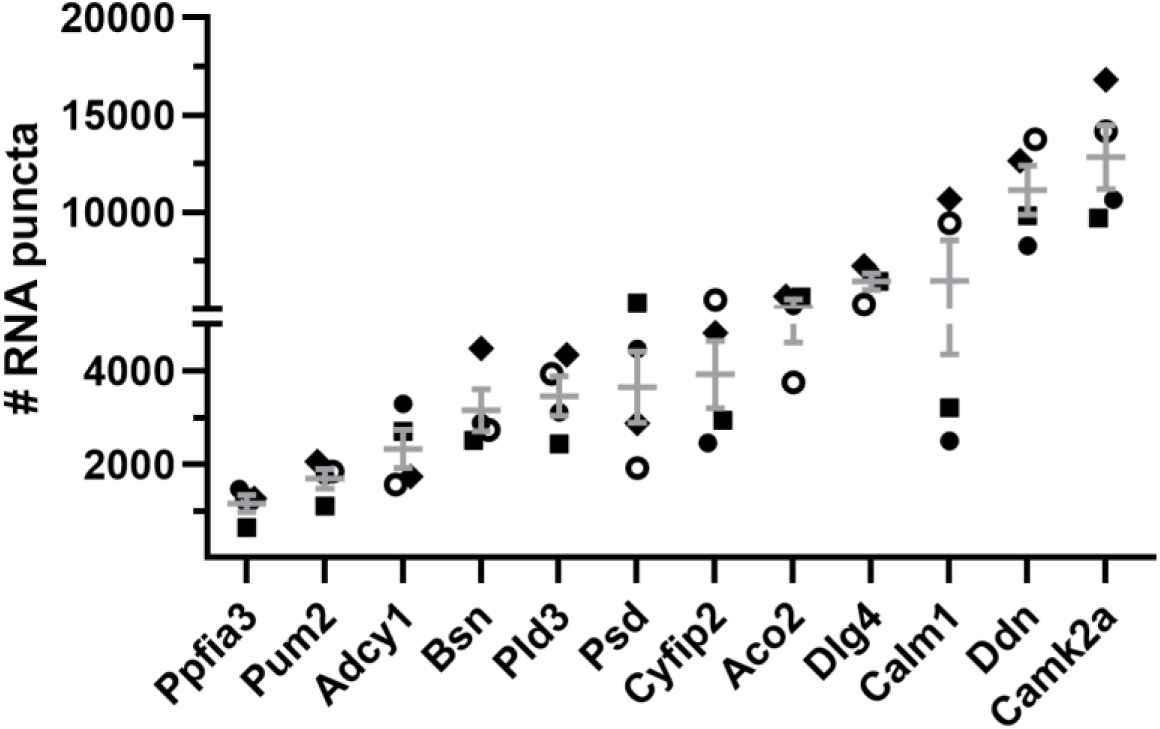
Abundance of each mRNA in the CA2 neuropil. Each symbol represents data from a biological replicate (N=4 mice). Error bars indicates SEM.

**Supplemental Fig. 5 (Refers to Fig. 3A).**
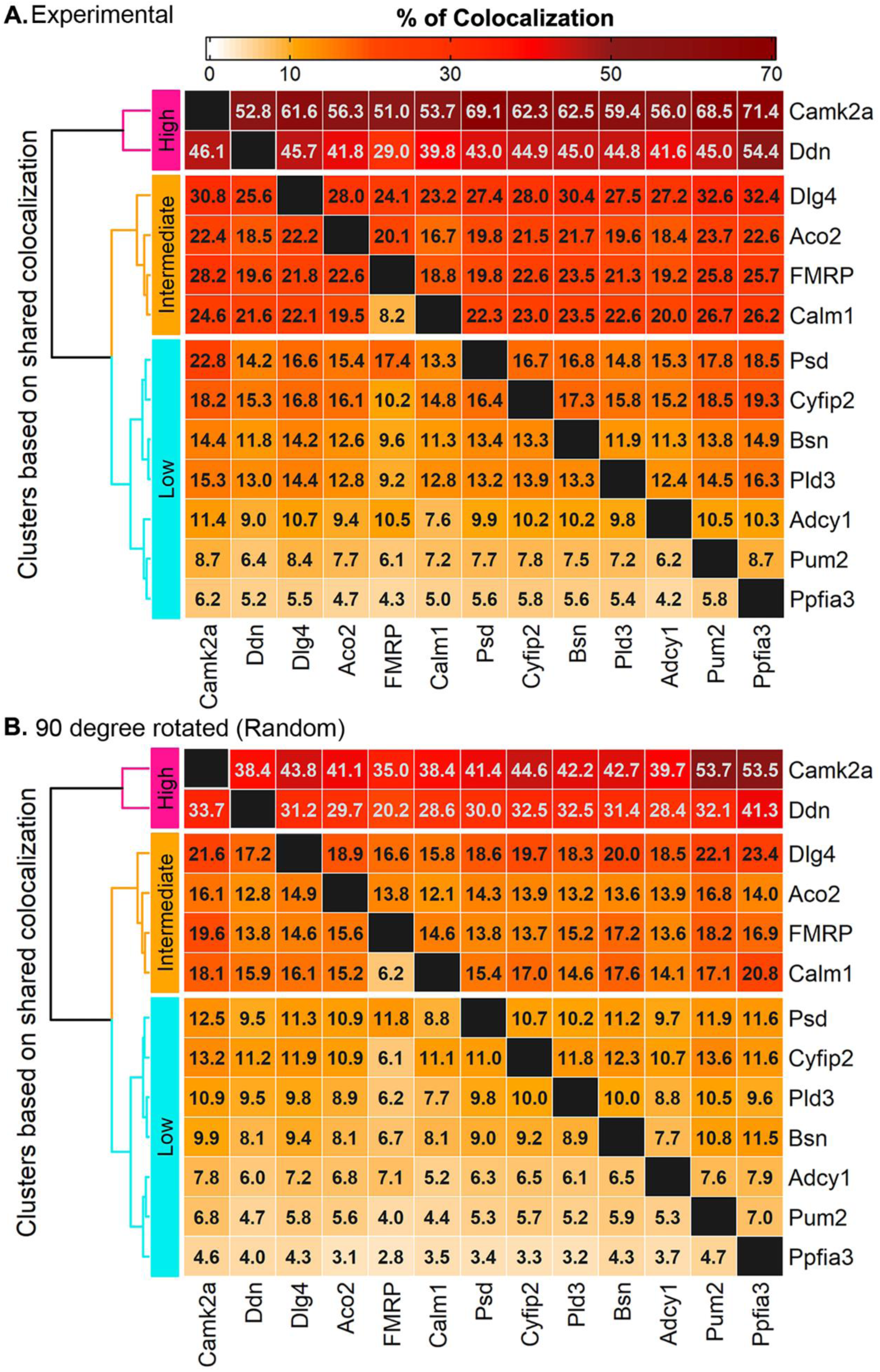
Heatmaps showing the total average pairwise colocalization of mRNAs in properly registered (experimental) images **(A)** and in rotated (random) images **(B)**. The percentage values in each column are calculated by dividing the number of column mRNAs colocalizing with each row mRNA by the total number of column mRNAs, i.e. 5.2% of *Ddn* colocalizes with *Ppfia3*, whereas 54.4% of *Ppfia3* colocalizes with *Ddn* before random colocalization subtraction (see Fig. 4A). Values are the average of N=4 mice (four 52 X 52 µm^2^ images averaged per mouse).

**Supplemental Fig. 6:**
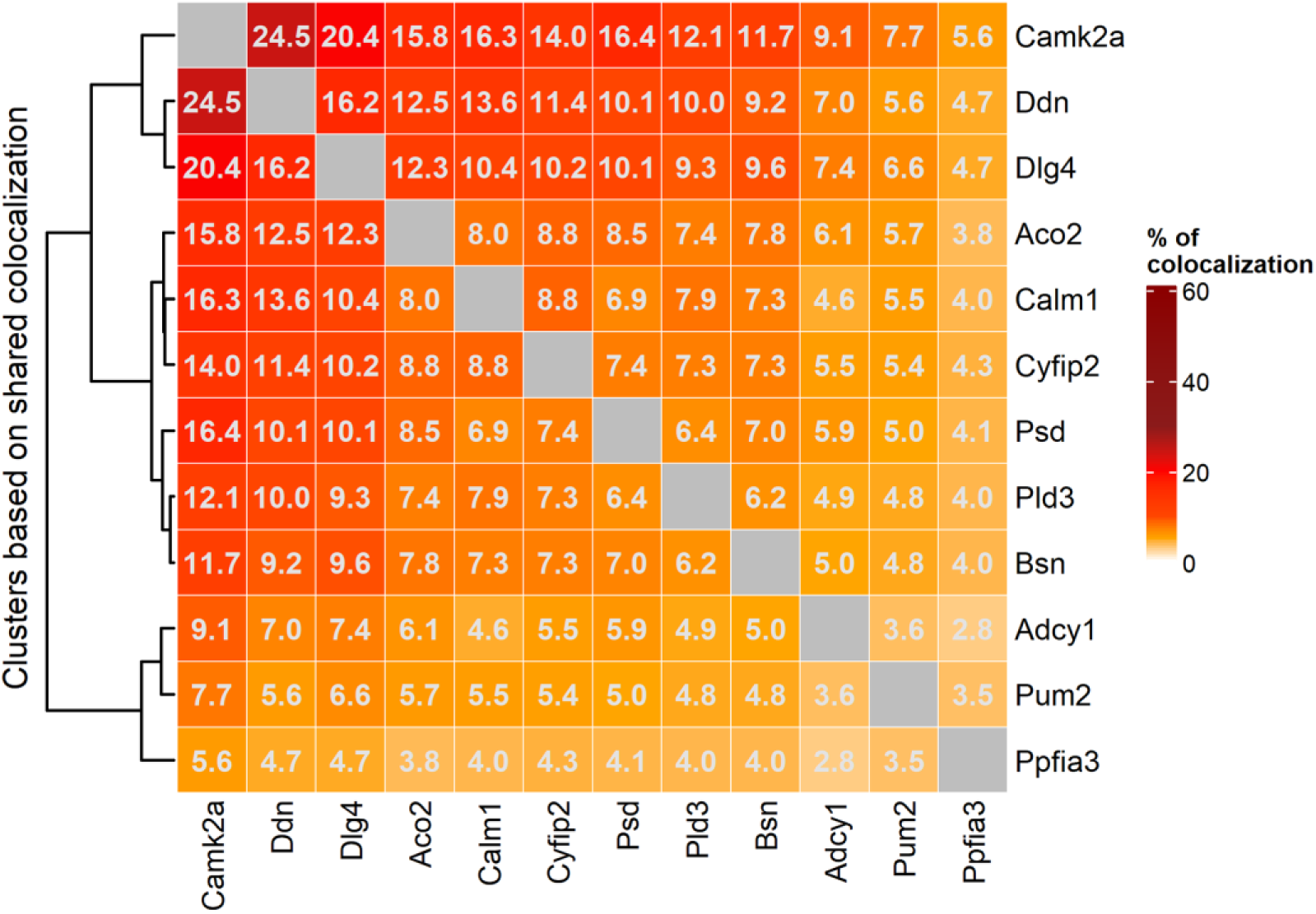
Pairwise colocalization of neuropil localized mRNAs analyzed as in Batish et al. (Batish et al., 2012). For each pair of comparisons, the number of overlapping RNA puncta between two channels was divided by the combined count of the two RNAs being compared and expressed as a percentage (average of N=4 mice). Hierarchical clustering of the data revealed a very similar pattern (as shown in Fig 3A) showing that every RNA is colocalized more with highly abundant RNAs (*Camk2a, Ddn, Dlg4*) and show fewer instances of colocalization with RNAs that are of lower abundance (*Pum2, Ppfia3*). RNAs in intermediary clusters also show a similar trend although their specific orders are more variable compared to Fig 3A.

**Supplemental Fig. 7 (Refers to Fig. 3C).**
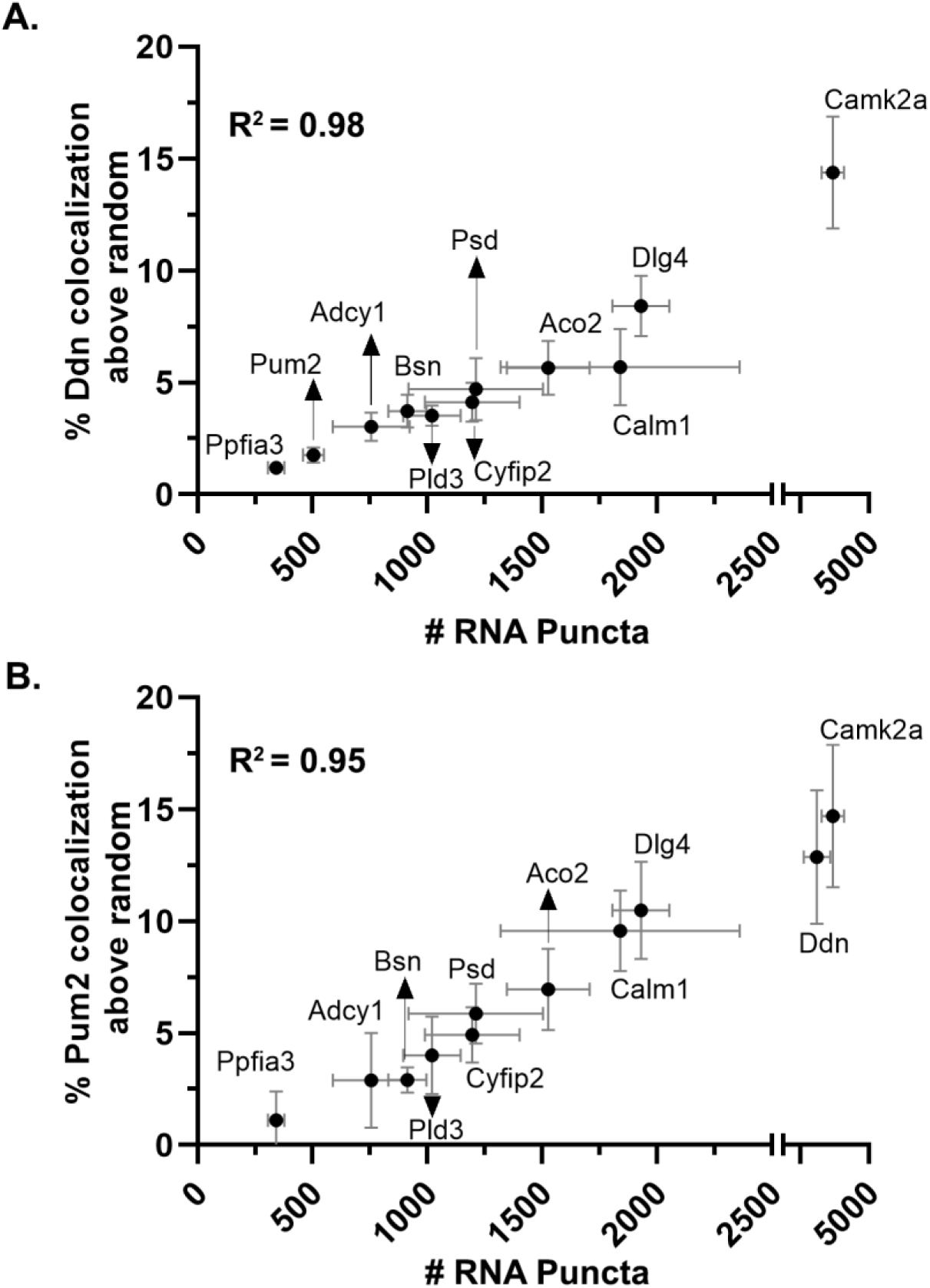
The positive correlation between pairwise colocalization and mRNA abundance exists regardless of expression. *Ddn* (A) and *Pum2* (B) exhibit high and low abundance, respectively, in CA2 neuropil. However, these mRNAs display a consistent positive correlation between % colocalization (random colocalization subtracted) and the abundance of the 11 paired mRNAs (*Ddn* R^2^ = 0.98 and *Pum2* R^2^ = 0.95). (N=4 mice. Error bars indicate SEM.)

**Supplemental Fig. 8 (Refers to Fig. 3D).**
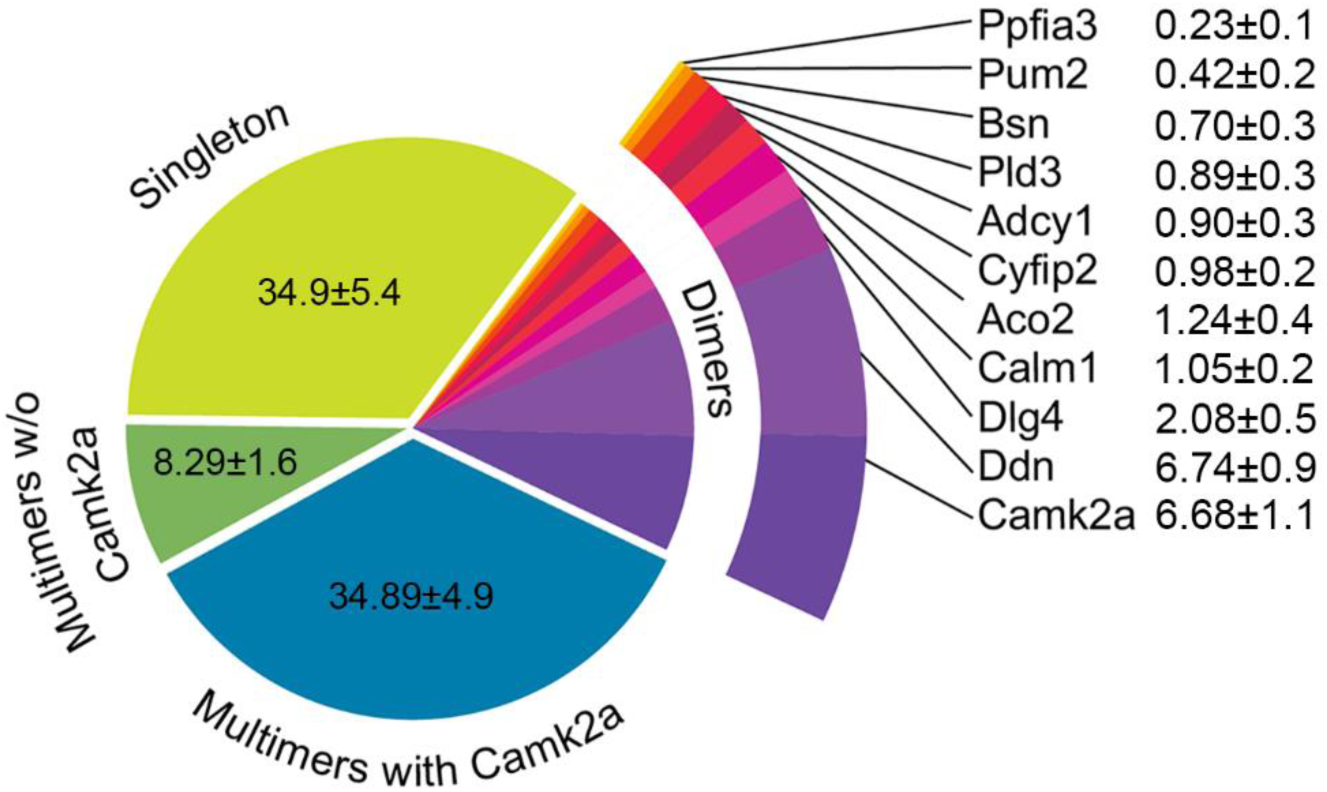
Pie chart of *Psd* mRNA composition that was observed due to random overlap of mRNA fluorescent puncta. *Psd* image was rotated 90 degrees and colocalization of *Psd* with other eleven mRNAs combined were quantified and averaged from the same four 52X52 µm^2^ ROIs per animal as done for the registered experimental images. Individual animal averages were then averaged across N=4 mice. 65.1 ± 5.4% of *Psd* mRNA puncta (vs. 91.86 ± 1.8% in properly registered images) overlap randomly with at least one other mRNA that include dimers (*Psd* with only one other mRNA) or multimers (*Psd* with at least two other mRNAs). Consistent with the pairwise colocalization data where the extent of colocalization scales with mRNA abundance, the percentage of randomly colocalized dimers increases as mRNA abundance increases. However, the percentage of random dimers is equal to or greater than the percentage of dimers from properly registered images, with the exception of *Psd/Camk2a* dimers that are present at lower percentage than experimental (random *Psd/Camk2a* dimers 6.68 ± 1.1% versus properly registered *Psd/Camk2a* dimers 11.58 ± 3.1%). Random *Psd*-multimers with *Camk2a* (34.9 ± 5.0%) and without *Camk2a* (8.29 ± 1.6%) are appreciably lower than experimental images (57.91 ± 5.3% and 12.34 ± 2.9%, respectively), indicating mul- timer populations dominate colocalized Psd RNA puncta compositions in our data.

**Supplemental Fig. 9 (refers to Fig. 3D).**
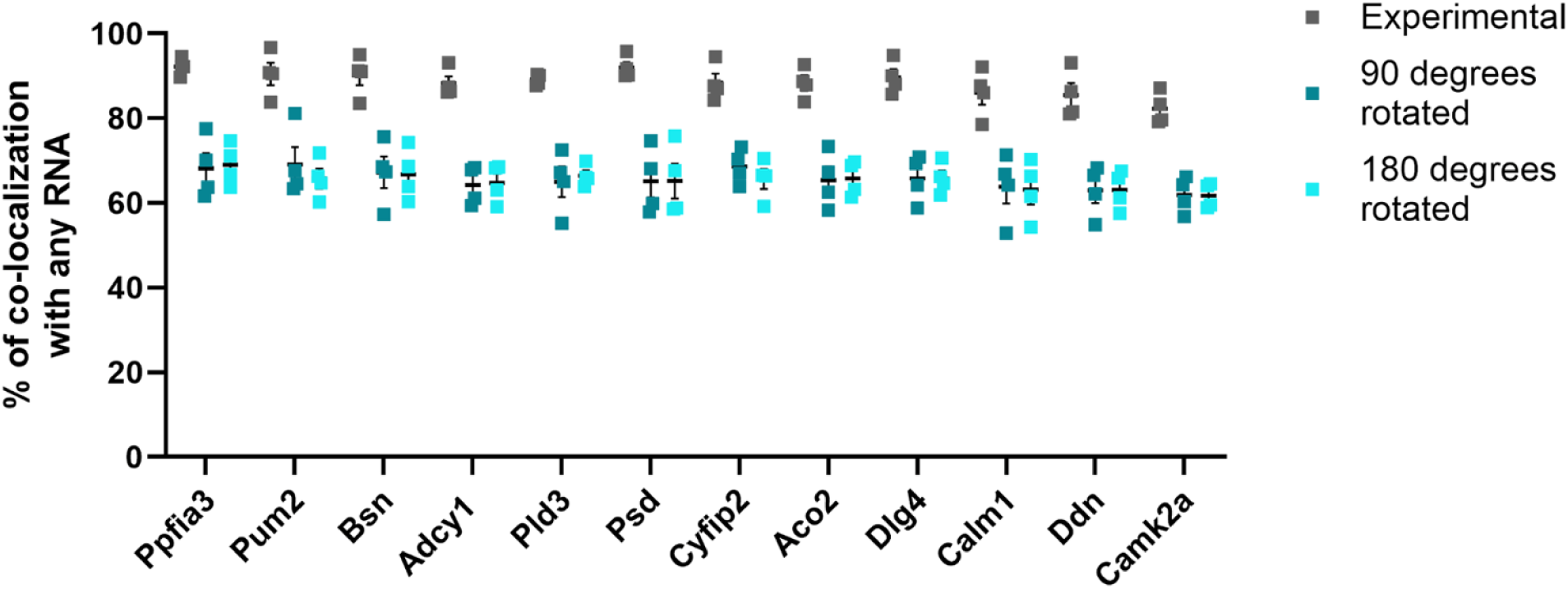
The majority of neuropil localized mRNAs spatially interact with at least one other RNA. Total % colocalization of each mRNA with any of the other 11 mRNAs from properly registered experimental images and 90 degree as well as 180-degree rotated images. Each symbol represents a biological replicate. % colocalization from experimental images were significantly higher compared to that in 90-degree rotated images (unpaired two sample multiple t-test with Welch’s correction, p<0.01 for every pair) and 180-degree rotated images (unpaired two sample multiple t-test with Welch’s correction, p<0.01 for every pair). N=4 mice. Error bars indicates SEM.

**Supplemental Fig. 10 (Refers to Fig. 3).**
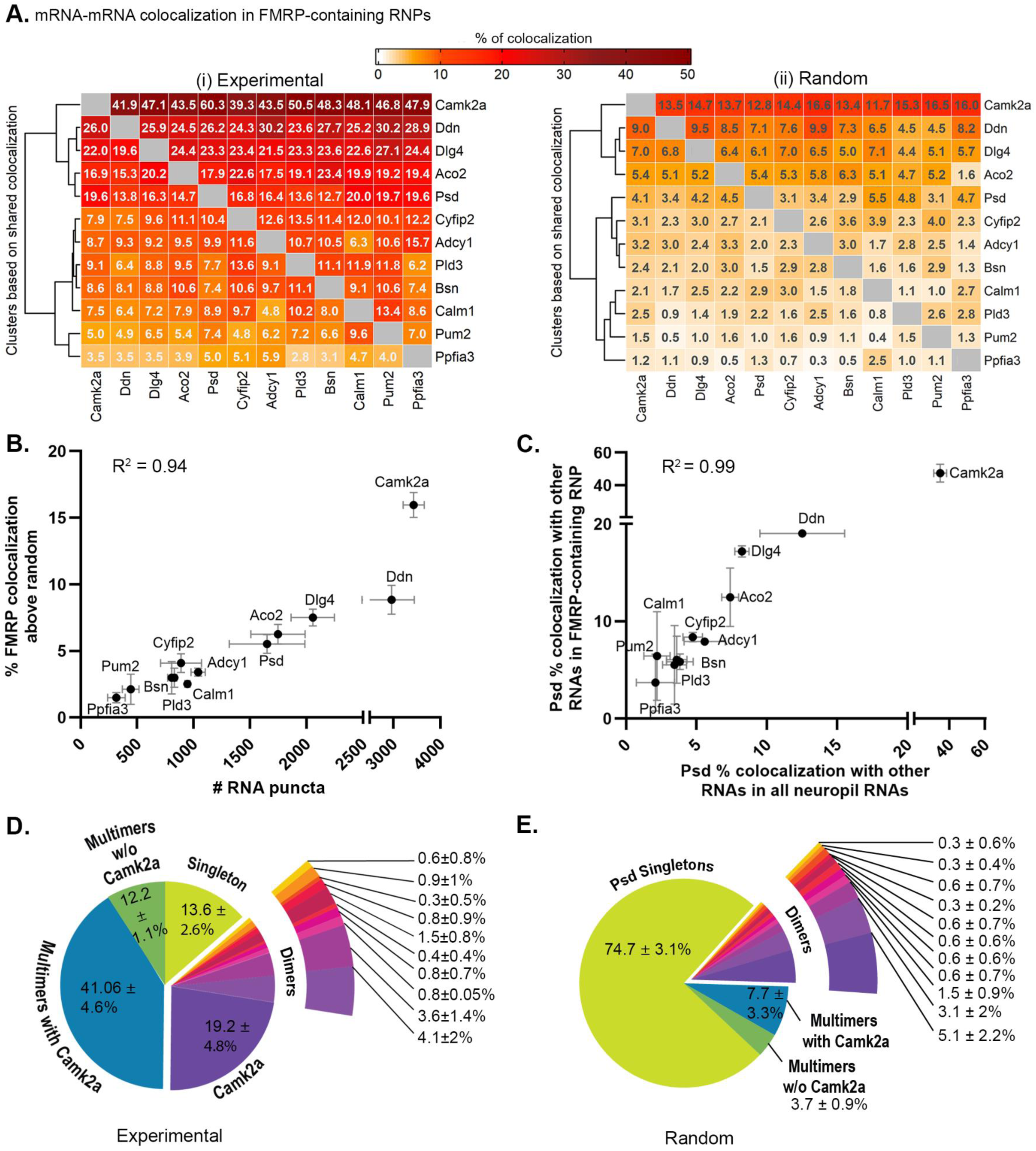
Colocalization of mRNAs within FMRP-containing RNPs. **A.** Heatmaps of experimental (Ai. left) and random (Aii. right, 90 degree rotated) % colocalization of each mRNA pair within FMRP containing RNP. Percentage was calculated by dividing the number of overlapping puncta by the total number of the column mRNA puncta. **B.** Correlation plot of the percent FMRP colocalized with each mRNA (random colocalization subtracted) and mRNA abundance (R^2^ = 0.94). **C.** Correlation plot of pairwise *Psd* colocalization percentage with other RNAs within all neuropil RNA puncta (X-axis) and FMRP-containing RNPs (Y-axis). **D.** FMRP-containing *Psd* RNP compositions in experimental images. **E.** FMRP-containing *Psd* RNP compositions in random 90-degree rotated images. (N=2 mice).

**Supplemental Fig. 11 (refers to Fig. 4).**
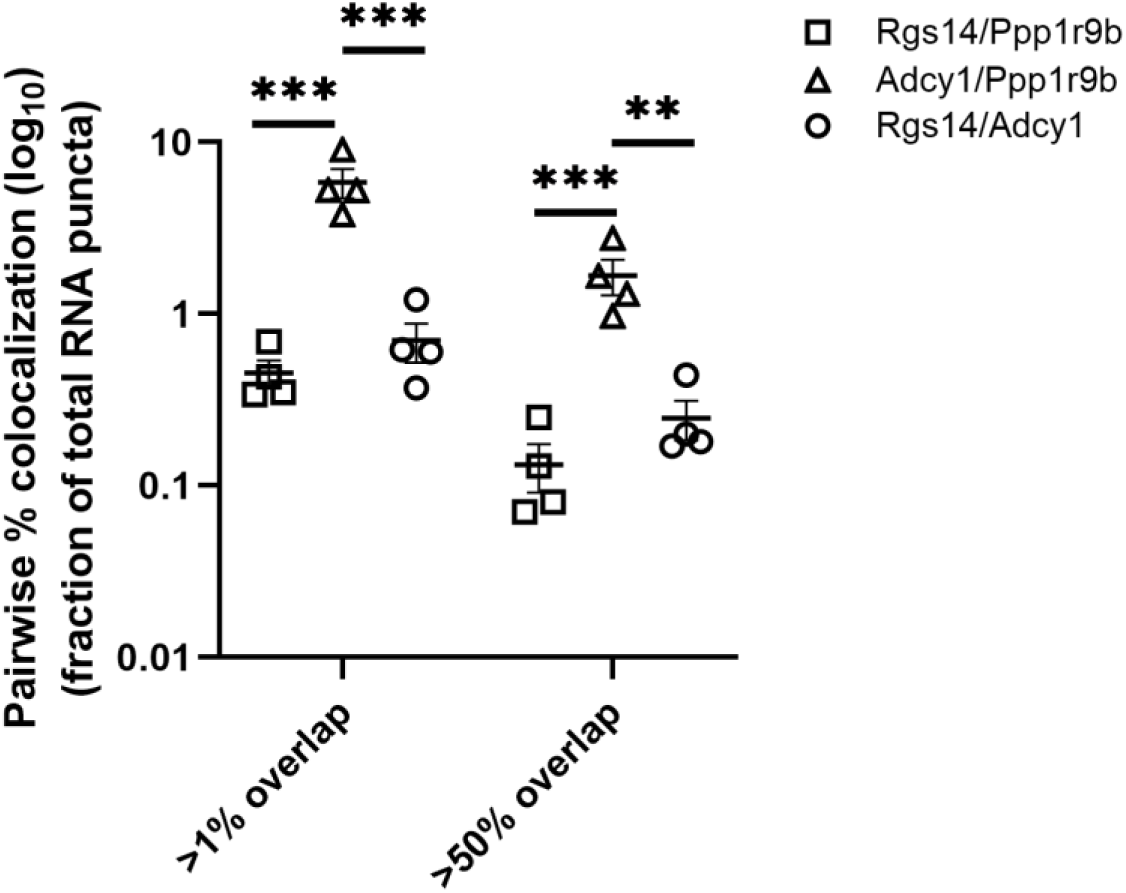
Correlation of mRNA pairwise % colocalization with mRNA abundance is consistent across stringent definitions of colocalization. mRNA pairwise colocalization is expressed as a percentage of the combined total mRNA puncta count in DG. Two highly abundant mRNAs *Adcy1* and *Ppp1r9b* are colocalized (5.84 ± 1.13%) significantly higher than *Adcy1/Rgs14* (0.70 ± 0.18%) and *Rgs14/Ppp1r9b* (0.45 ± 0.08%) mRNA pairs when colocalization is defined as >1% overlap (significant effect of mRNA pair, RM ANOVA: F = 49.26, p = 0.0002, N=4 mice) and this effect is significant for both comparisons (*Adcy1/Ppp1r9b* vs *Adcy1/Rgs14*: p = 0.0006; *Adcy1/Ppp1r9b* vs *Rgs14/Ppp1r9b*: p = 0.0002, Tukey’s post hoc tests). This difference in colocalization between mRNA pairs was recapitulated when colocalization was defined as >50% overlap between channels (significant effect of mRNA pair, RM ANOVA, F = 28.80, p = 0.0008, N=4 mice) and this was also significant for both comparisons (*Adcy1/Ppp1r9b* vs *Adcy1/Rgs14*: p = 0.0040; *Adcy1/Ppp1r9b* vs *Rgs14/Ppp1r9b*: p = 0.0008, Tukey’s post hoc tests). Stats were run on the transformed (log_10_) values as plotted to meet the normality assumption. Tukey’s post hoc tests reported on the plot.

**Supplemental Fig. 12 (refers to Fig. 4).**
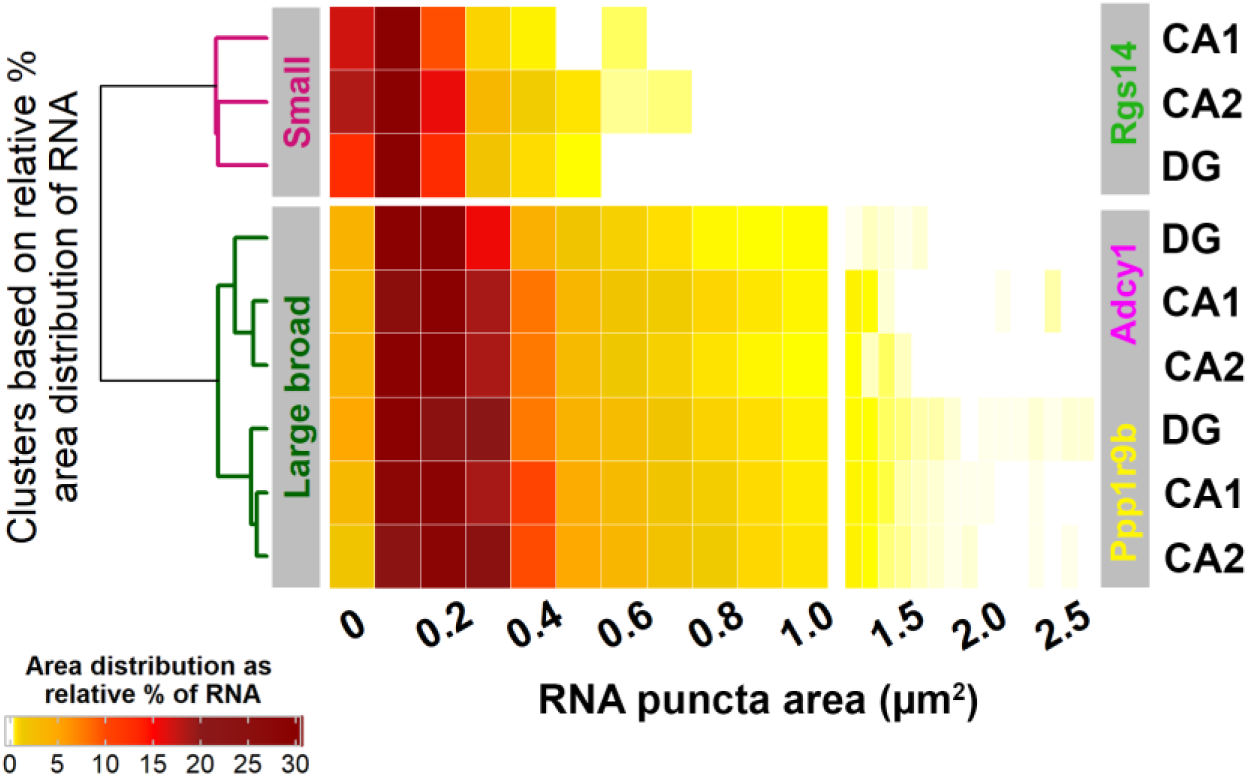
*Rgs14*, *Adcy1* and *Ppp1r9b* are variable in fluorescent puncta areas across mRNAs but not cell types. Hierarchical clustering of relative percent area distribution of *Rgs14*, *Adcy1* and *Ppp1r9b* in CA2, CA1 and DG of adult mouse hippocampus. Median puncta areas were not significantly different across cell types for each mRNA although heterogeneity in area distribution across mRNAs was observed similar to the HiPlex data. *Rgs14* median puncta area CA1: 0.10 ± 0.01 µm^2^, CA2: 0.10 ± 0.01 um^2^, DG: 0.10 ± 0.02 um^2^ (no effect of cell-type, RM ANOVA, F = 0.026, p = 0.9744, N=4 mice). *Adcy1* median puncta area CA1: 0.22 ± 0.03 µm^2^, CA2: 0.22 ± 0.03 µm^2^, DG: 0.19 ± 0.02 µm^2^ (no effect of cell type, RM ANOVA: F = 0.5406, p = 0.6083, N=4 mice). *Ppp1r9b* median puncta area CA1: 0.22 ± 0.01 µm^2^, CA2: 0.24 ± 0.02 µm^2^, DG: 0.22 ± 0.03 µm^2^ (no-effect of cell type, RM ANOVA, F = 0.5507, p = 0.6032, N=4 mice). Since *Adcy1* and *Ppp1r9b* median puncta area data were more likely to be a lognormal distribution, we repeated the RM ANOVA on log10 transformed values, which did not change the statistical result.

## TABLES

**Supp. Table 1 (Refers to Fig. 2 and 3).**
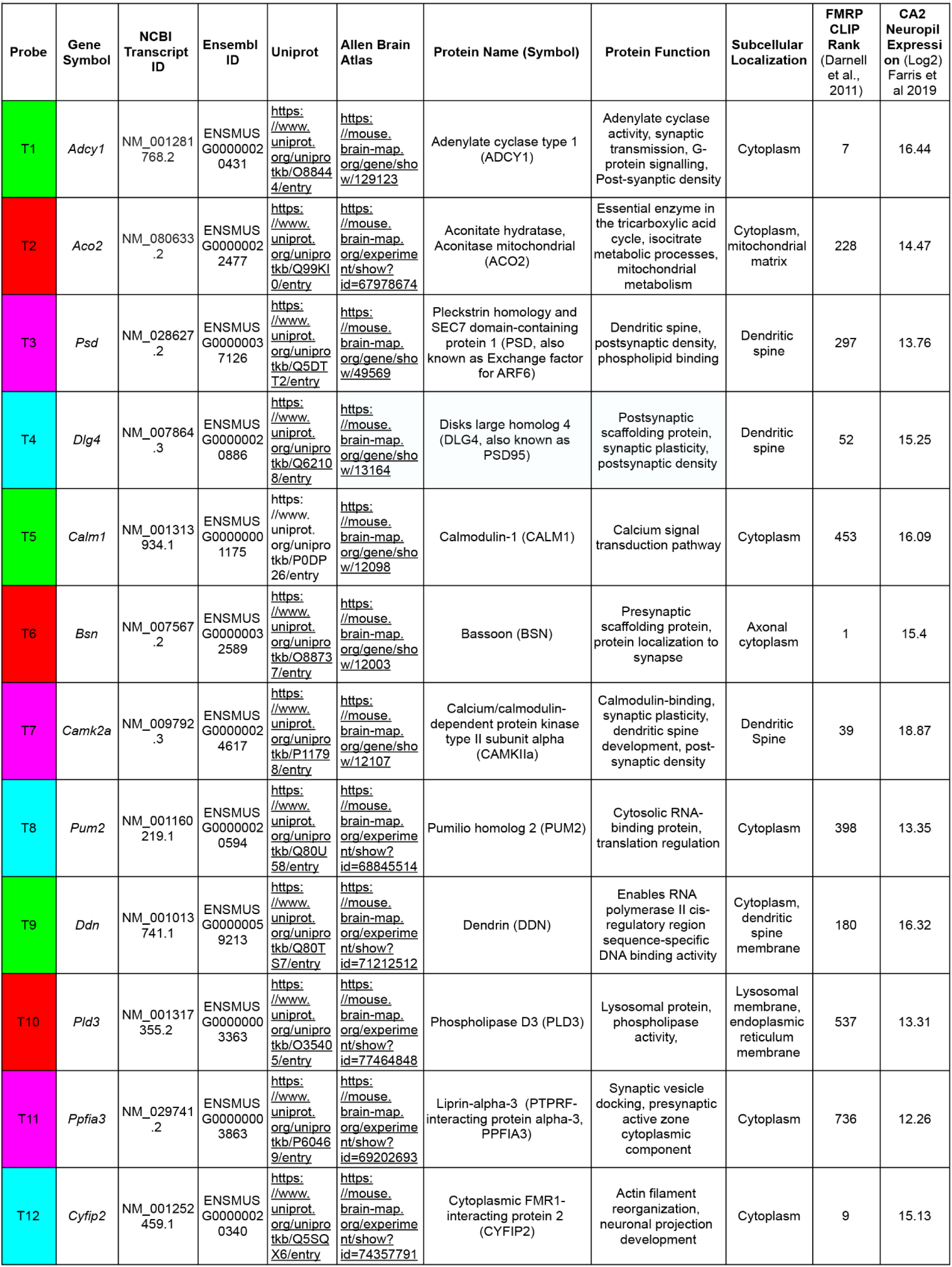
HiPlex probe information. See Supplemental file for full datasheet.

**Supplemental Table 2:**
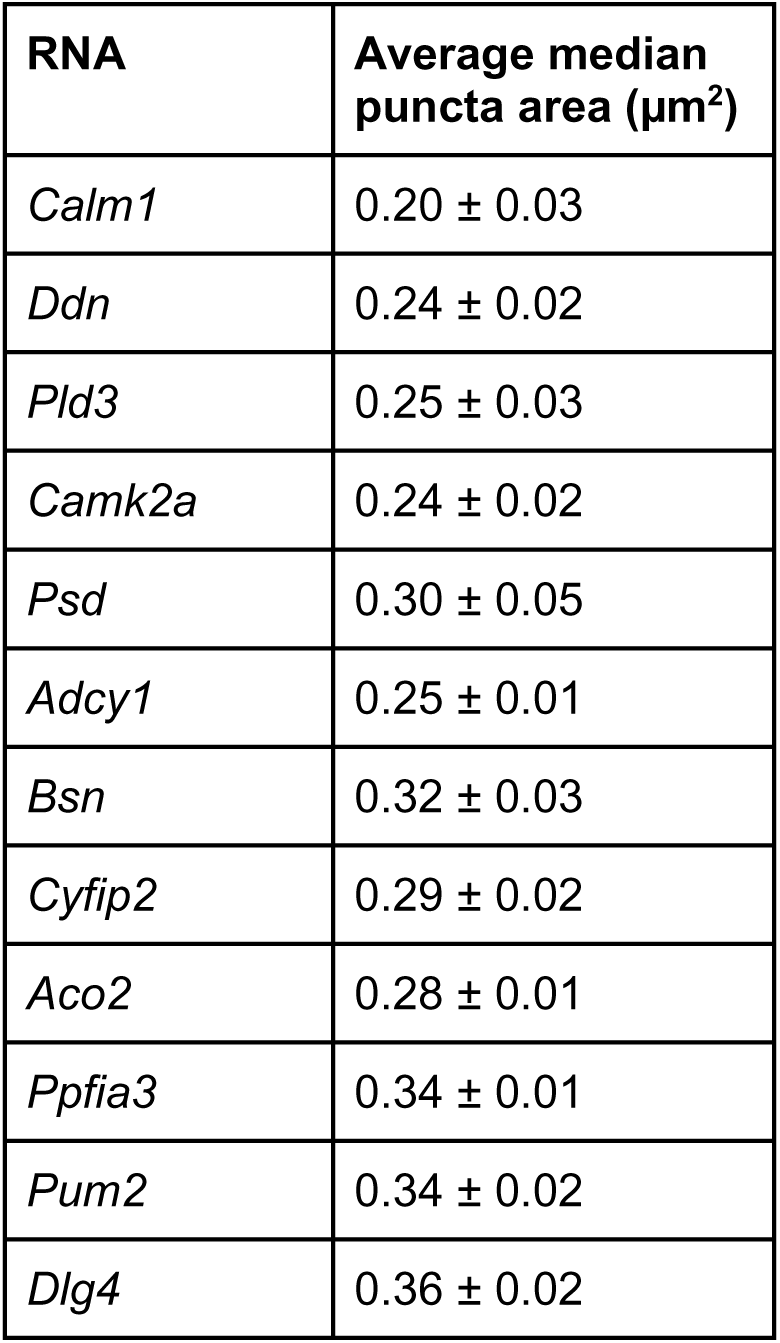
Average median size of twelve mRNAs visualized in HiPlex smFISH (N=4 mice).

## REFERENCES

Ahn, H., Durang, X., Shim, J. Y., Park, G., Jeon, J., & Park, H. Y. (2023). Statistical modeling of mRNP transport in dendrites: A comparative analysis of β-actin and Arc mRNP dynamics. *Traffic (Copenhagen*, Denmark), 24(11), 522–532. 10.1111/tra.12913

Ainsley, J. A., Drane, L., Jacobs, J., Kittelberger, K. A., & Reijmers, L. G. (2014). Functionally diverse dendritic mRNAs rapidly associate with ribosomes following a novel experience. Nature Communications, 5(1), 4510. 10.1038/ncomms5510

Antar, L. N., Dictenberg, J. B., Plociniak, M., Afroz, R., & Bassell, G. J. (2005). Localization of FMRP-associated mRNA granules and requirement of microtubules for activity-dependent trafficking in hippocampal neurons. *Genes*, Brain and Behavior, 4(6), 350–359. 10.1111/j.1601-183X.2005.00128.x

Batish, M., van den Bogaard, P., Kramer, F. R., & Tyagi, S. (2012). Neuronal mRNAs travel singly into dendrites. Proceedings of the National Academy of Sciences, 109(12), 4645–4650. 10.1073/pnas.1111226109

Bauer, K. E., Bargenda, N., Schieweck, R., Illig, C., Segura, I., Harner, M., & Kiebler, M. A. (2022). RNA supply drives physiological granule assembly in neurons. Nature Communications, 13(1), 2781. 10.1038/s41467-022-30067-3

Bauer, K. E., de Queiroz, B. R., Kiebler, M. A., & Besse, F. (2023). RNA granules in neuronal plasticity and disease. Trends in Neurosciences, 46(7), 525–538. 10.1016/j.tins.2023.04.004

Bauer, K. E., Segura, I., Gaspar, I., Scheuss, V., Illig, C., Ammer, G., Hutten, S., Basyuk, E., Fernández-Moya, S. M., Ehses, J., Bertrand, E., & Kiebler, M. A. (2019). Live cell imaging reveals 3′-UTR dependent mRNA sorting to synapses. Nature Communications, 10(1), 3178. 10.1038/s41467-01911123-x

Bergmann, C., Mousaei, K., Rizzoli, S. O., & Tchumatchenko, T. (2025). How energy determines spatial localisation and copy number of molecules in neurons. Nature Communications, 16(1), 1424. 10.1038/s41467-025-56640-0

Boeynaems, S., Holehouse, A. S., Weinhardt, V., Kovacs, D., Van Lindt, J., Larabell, C., Van Den Bosch, L., Das, R., Tompa, P. S., Pappu, R. V., & Gitler, A. D. (2019). Spontaneous driving forces give rise to protein−RNA condensates with coexisting phases and complex material properties. Proceedings of the National Academy of Sciences, 116(16), 7889–7898. 10.1073/pnas.1821038116

Bolte, S., & Cordelières, F. P. (2006). A guided tour into subcellular colocalization analysis in light microscopy. Journal of Microscopy, 224(Pt 3), 213–232. 10.1111/j.1365-2818.2006.01706.x

Bradshaw, K. D., Emptage, N. J., & Bliss, T. V. P. (2003). A role for dendritic protein synthesis in hippocampal late LTP. European Journal of Neuroscience, 18(11), 3150–3152. 10.1111/j.1460-9568.2003.03054.x

Buxbaum, A. R., Haimovich, G., & Singer, R. H. (2015). In the right place at the right time: Visualizing and understanding mRNA localization. Nature Reviews. Molecular Cell Biology, 16(2), 95–109. 10.1038/nrm3918

Cajigas, I. J., Tushev, G., Will, T. J., Dieck, S. tom, Fuerst, N., & Schuman, E. M. (2012). The Local Transcriptome in the Synaptic Neuropil Revealed by Deep Sequencing and High-Resolution Imaging. Neuron, 74(3), 453–466. 10.1016/j.neuron.2012.02.036

Carson, J. H., Gao, Y., Tatavarty, V., Levin, M. K., Korza, G., Francone, V. P., Kosturko, L. D., Maggipinto, M. J., & Barbarese, E. (2008). Multiplexed RNA trafficking in oligodendrocytes and neurons. Biochimica et Biophysica Acta (BBA) - Gene Regulatory Mechanisms, 1779(8), 453–458. 10.1016/j.bbagrm.2008.04.002

Cirillo, D., Agostini, F., Klus, P., Marchese, D., Rodriguez, S., Bolognesi, B., & Tartaglia, G. G. (2013). Neurodegenerative diseases: Quantitative predictions of protein–RNA interactions. RNA, 19(2), 129–140. 10.1261/rna.034777.112

Cougot, N., Bhattacharyya, S. N., Tapia-Arancibia, L., Bordonné, R., Filipowicz, W., Bertrand, E., & Rage, F. (2008). Dendrites of Mammalian Neurons Contain Specialized P-Body-Like Structures That Respond to Neuronal Activation. The Journal of Neuroscience, 28(51), 13793–13804. 10.1523/JNEUROSCI.4155-08.2008

Darnell, J. C., Van Driesche, S. J., Zhang, C., Hung, K. Y. S., Mele, A., Fraser, C. E., Stone, E. F., Chen, C., Fak, J. J., Chi, S. W., Licatalosi, D. D., Richter, J. D., & Darnell, R. B. (2011). FMRP Stalls Ribosomal Translocation on mRNAs Linked to Synaptic Function and Autism. Cell, 146(2), 247–261. 10.1016/j.cell.2011.06.013

Dictenberg, J. B., Swanger, S. A., Antar, L. N., Singer, R. H., & Bassell, G. J. (2008). A Direct Role for FMRP in Activity-Dependent Dendritic mRNA Transport Links Filopodial-Spine Morphogenesis to Fragile X Syndrome. Developmental Cell, 14(6), 926–939. 10.1016/j.devcel.2008.04.003

Donlin-Asp, P. G., Polisseni, C., Klimek, R., Heckel, A., & Schuman, E. M. (2021). Differential regulation of local mRNA dynamics and translation following long-term potentiation and depression. Proceedings of the National Academy of Sciences, 118(13), e2017578118. 10.1073/pnas.2017578118

Dunn, K. W., Kamocka, M. M., & McDonald, J. H. (2011). A practical guide to evaluating colocalization in biological microscopy. American Journal of Physiology - Cell Physiology, 300(4), C723–C742. 10.1152/ajpcell.00462.2010

Elvira, G., Wasiak, S., Blandford, V., Tong, X.-K., Serrano, A., Fan, X., Sánchez-Carbente, M. del R., Servant, F., Bell, A. W., Boismenu, D., Lacaille, J.-C., McPherson, P. S., DesGroseillers, L., & Sossin, W. S. (2006). Characterization of an RNA Granule from Developing Brain *. Molecular & Cellular Proteomics, 5(4), 635–651. 10.1074/mcp.M500255-MCP200

Evans, P. R., Gerber, K. J., Dammer, E. B., Duong, D. M., Goswami, D., Lustberg, D. J., Zou, J., Yang, J. J., Dudek, S. M., Griffin, P. R., Seyfried, N. T., & Hepler, J. R. (2018). Interactome Analysis Reveals Regulator of G Protein Signaling 14 (RGS14) is a Novel Calcium/Calmodulin (Ca2+/CaM) and CaM Kinase II (CaMKII) Binding Partner. Journal of Proteome Research, 17(4), 1700–1711. 10.1021/acs.jproteome.8b00027

Farris, S., Lewandowski, G., Cox, C. D., & Steward, O. (2014). Selective Localization of Arc mRNA in Dendrites Involves Activity- and Translation-Dependent mRNA Degradation. The Journal of Neuroscience, 34(13), 4481–4493. 10.1523/JNEUROSCI.4944-13.2014

Farris, S., Ward, J. M., Carstens, K. E., Samadi, M., Wang, Y., & Dudek, S. M. (2019). Hippocampal Subregions Express Distinct Dendritic Transcriptomes that Reveal Differences in Mitochondrial Function in CA2. Cell Reports, 29(2), 522–539.e6. 10.1016/j.celrep.2019.08.093

Fatimy, R. E., Davidovic, L., Tremblay, S., Jaglin, X., Dury, A., Robert, C., Koninck, P. D., & Khandjian, E. W. (2016). Tracking the Fragile X Mental Retardation Protein in a Highly Ordered Neuronal RiboNucleo-Particles Population: A Link between Stalled Polyribosomes and RNA Granules. PLOS Genetics, 12(7), e1006192. 10.1371/journal.pgen.1006192

Fernandez-Moya, S. M., Bauer, K. E., & Kiebler, M. A. (2014). Meet the players: Local translation at the synapse. Frontiers in Molecular Neuroscience, 7, 84. 10.3389/fnmol.2014.00084

Fernandopulle, M. S., Lippincott-Schwartz, J., & Ward, M. E. (2021). RNA transport and local translation in neurodevelopmental and neurodegenerative disease. Nature Neuroscience, 24(5), 622–632. 10.1038/s41593-020-00785-2

Ford, L., Ling, E., Kandel, E. R., & Fioriti, L. (2019). CPEB3 inhibits translation of mRNA targets by localizing them to P bodies. Proceedings of the National Academy of Sciences, 116(36), 18078–18087. 10.1073/pnas.1815275116

Fritzsche, R., Karra, D., Bennett, K. L., Ang, F. yee, Heraud-Farlow, J. E., Tolino, M., Doyle, M., Bauer, K. E., Thomas, S., Planyavsky, M., Arn, E., Bakosova, A., Jungwirth, K., Hörmann, A., Palfi, Z., Sandholzer, J., Schwarz, M., Macchi, P., Colinge, J., … Kiebler, M. A. (2013). Interactome of Two Diverse RNA Granules Links mRNA Localization to Translational Repression in Neurons. Cell Reports, 5(6), 1749–1762. 10.1016/j.celrep.2013.11.023

Gao, Y., Tatavarty, V., Korza, G., Levin, M. K., & Carson, J. H. (2008). Multiplexed Dendritic Targeting of α Calcium Calmodulin-dependent Protein Kinase II, Neurogranin, and Activity-regulated Cytoskeleton-associated Protein RNAs by the A2 Pathway. Molecular Biology of the Cell, 19(5), 2311–2327. 10.1091/mbc.e07-09-0914

Garcia-Jove Navarro, M., Kashida, S., Chouaib, R., Souquere, S., Pierron, G., Weil, D., & Gueroui, Z. (2019). RNA is a critical element for the sizing and the composition of phase-separated RNA–protein condensates. Nature Communications, 10(1), 3230. 10.1038/s41467-019-11241-6

Glock, C., Biever, A., Tushev, G., Bartnik, I., Nassim-Assir, B., Dieck, S. tom, & Schuman, E. M. (2020). The mRNA translation landscape in the synaptic neuropil (p. 2020.06.09.141960). bioRxiv. 10.1101/2020.06.09.141960

Hafner, A.-S., Donlin-Asp, P. G., Leitch, B., Herzog, E., & Schuman, E. M. (2019). Local protein synthesis is a ubiquitous feature of neuronal pre- and postsynaptic compartments. Science. 10.1126/science.aau3644

Hale, C. R., Sawicka, K., Mora, K., Fak, J. J., Kang, J. J., Cutrim, P., Cialowicz, K., Carroll, T. S., & Darnell, R. B. (2021). FMRP regulates mRNAs encoding distinct functions in the cell body and dendrites of CA1 pyramidal neurons. eLife, 10, e71892. 10.7554/eLife.71892

Heraud-Farlow, J. E., Sharangdhar, T., Li, X., Pfeifer, P., Tauber, S., Orozco, D., Hörmann, A., Thomas, S., Bakosova, A., Farlow, A. R., Edbauer, D., Lipshitz, H. D., Morris, Q. D., Bilban, M., Doyle, M., & Kiebler, M. A. (2013). Staufen2 Regulates Neuronal Target RNAs. Cell Reports, 5(6), 1511–1518. 10.1016/j.celrep.2013.11.039

Huang, Y.-S., Carson, J. H., Barbarese, E., & Richter, J. D. (2003). Facilitation of dendritic mRNA transport by CPEB. Genes & Development, 17(5), 638–653. 10.1101/gad.1053003

Huber, K. M., Gallagher, S. M., Warren, S. T., & Bear, M. F. (2002). Altered synaptic plasticity in a mouse model of fragile X mental retardation. Proceedings of the National Academy of Sciences, 99(11), 7746– 7750. 10.1073/pnas.122205699

Huber, K. M., Kayser, M. S., & Bear, M. F. (2000). Role for rapid dendritic protein synthesis in hippocampal mGluR-dependent long-term depression. *Science (New York*, N.Y.), 288(5469), 1254–1257. 10.1126/science.288.5469.1254

Jiang, Y.-H., & Ehlers, M. D. (2013). Modeling autism by SHANK gene mutations in mice. Neuron, 78(1), 8–27. 10.1016/j.neuron.2013.03.016

Kanai, Y., Dohmae, N., & Hirokawa, N. (2004). Kinesin Transports RNA: Isolation and Characterization of an RNA-Transporting Granule. Neuron, 43(4), 513–525. 10.1016/j.neuron.2004.07.022

Kharod, S. C., Hwang, D., Castillo, P. E., & Yoon, Y. J. (2023). Regulation of FMRP granule structure and function through phosphorylation (p. 2023.03.15.532613). bioRxiv. 10.1101/2023.03.15.532613

Kiebler, M. A., & Bassell, G. J. (2006). Neuronal RNA Granules: Movers and Makers. Neuron, 51(6), 685–690. 10.1016/j.neuron.2006.08.021

Kiebler, M. A., & Bauer, K. E. (2024). RNA granules in flux: Dynamics to balance physiology and pathology | Nature Reviews Neuroscience. Nature Reviewes Neuroscience, 711–725.

Knowles, R. B., Sabry, J. H., Martone, M. E., Deerinck, T. J., Ellisman, M. H., Bassell, G. J., & Kosik, K. S. (1996). Translocation of RNA Granules in Living Neurons. Journal of Neuroscience, 16(24), 7812– 7820. 10.1523/JNEUROSCI.16-24-07812.1996

Krichevsky, A. M., & Kosik, K. S. (2001). Neuronal RNA granules: A link between RNA localization and stimulation-dependent translation. Neuron, 32(4), 683–696. 10.1016/s0896-6273(01)00508-6

Lewis, Y. E., Moskovitz, A., Mutlak, M., Heineke, J., Caspi, L. H., & Kehat, I. (2018). Localization of transcripts, translation, and degradation for spatiotemporal sarcomere maintenance. Journal of Molecular and Cellular Cardiology, 116, 16–28. 10.1016/j.yjmcc.2018.01.012

Lipshitz, H. D., & Smibert, C. A. (2000). Mechanisms of RNA localization and translational regulation. Current Opinion in Genetics & Development, 10(5), 476–488. 10.1016/S0959-437X(00)00116-7

Maharana, S., Wang, J., Papadopoulos, D. K., Richter, D., Pozniakovsky, A., Poser, I., Bickle, M., Rizk, S., Guillén-Boixet, J., Franzmann, T. M., Jahnel, M., Marrone, L., Chang, Y.-T., Sterneckert, J., Tomancak, P., Hyman, A. A., & Alberti, S. (2018). RNA buffers the phase separation behavior of prion-like RNA binding proteins. Science, 360(6391), 918–921. 10.1126/science.aar7366

Martin, K. C., & Ephrussi, A. (2009). mRNA localization: Gene expression in the spatial dimension. Cell, 136(4), 719–730. 10.1016/j.cell.2009.01.044

McDonald, J. H., & Dunn, K. W. (2013). Statistical tests for measures of colocalization in biological microscopy. Journal of Microscopy, 252(3), 295–302. 10.1111/jmi.12093

Middleton, S. A., Eberwine, J., & Kim, J. (2019). Comprehensive catalog of dendritically localized mRNA isoforms from sub-cellular sequencing of single mouse neurons. BMC Biology, 17, 5. 10.1186/s12915-019-0630-z

Mikl, M., Vendra, G., & Kiebler, M. A. (2011). Independent localization of MAP2, CaMKIIα and β-actin RNAs in low copy numbers. EMBO Reports, 12(10), 1077–1084. 10.1038/embor.2011.149

Mitsumori, K., Takei, Y., & Hirokawa, N. (2017). Components of RNA granules affect their localization and dynamics in neuronal dendrites. Molecular Biology of the Cell, 28(11), 1412–1417. 10.1091/mbc.e16-07-0497

Miyashiro, K. Y., Beckel-Mitchener, A., Purk, T. P., Becker, K. G., Barret, T., Liu, L., Carbonetto, S., Weiler, I. J., Greenough, W. T., & Eberwine, J. (2003). RNA Cargoes Associating with FMRP Reveal Deficits in Cellular Functioning in Fmr1 Null Mice. Neuron, 37(3), 417–431. 10.1016/S0896-6273(03)00034-5

Monteiro, P., & Feng, G. (2017). SHANK proteins: Roles at the synapse and in autism spectrum disorder. Nature Reviews Neuroscience, 18(3), 147–157. 10.1038/nrn.2016.183

Moon, S. L., Morisaki, T., Khong, A., Lyon, K., Parker, R., & Stasevich, T. J. (2019). Multicolour single-molecule tracking of mRNA interactions with RNP granules. Nature Cell Biology, 21(2), 162–168. 10.1038/s41556-018-0263-4

Morris, C. W., Watkins, D. S., Shah, N. R., Pennington, T., Hens, B., Qi, G., Doud, E. H., Mosley, A. L., Atwood, B. K., & Baucum, A. J. (2023). Spinophilin Limits Metabotropic Glutamate Receptor 5 Scaffolding to the Postsynaptic Density and Cell Type Specifically Mediates Excessive Grooming. Biological Psychiatry, 93(11), 976–988. 10.1016/j.biopsych.2022.12.008

Nakayama, K., Ohashi, R., Shinoda, Y., Yamazaki, M., Abe, M., Fujikawa, A., Shigenobu, S., Futatsugi, A., Noda, M., Mikoshiba, K., Furuichi, T., Sakimura, K., & Shiina, N. (n.d.). RNG105/caprin1, an RNA granule protein for dendritic mRNA localization, is essential for long-term memory formation. eLife, 6, e29677. 10.7554/eLife.29677

Napoli, I., Mercaldo, V., Boyl, P. P., Eleuteri, B., Zalfa, F., De Rubeis, S., Di Marino, D., Mohr, E., Massimi, M., Falconi, M., Witke, W., Costa-Mattioli, M., Sonenberg, N., Achsel, T., & Bagni, C. (2008). The fragile X syndrome protein represses activity-dependent translation through CYFIP1, a new 4E-BP. Cell, 134(6), 1042–1054. 10.1016/j.cell.2008.07.031

Niepielko, M. G., Eagle, W. V. I., & Gavis, E. R. (2018). Stochastic Seeding Coupled with mRNA Self-Recruitment Generates Heterogeneous Drosophila Germ Granules. Current Biology: CB, 28(12), 1872–1881.e3. 10.1016/j.cub.2018.04.037

Ortiz, R., Georgieva, M. V., Gutiérrez, S., Pedraza, N., Fernández-Moya, S. M., & Gallego, C. (2017). Recruitment of Staufen2 Enhances Dendritic Localization of an Intron-Containing *CaMKIIα* mRNA. Cell Reports, 20(1), 13–20. 10.1016/j.celrep.2017.06.026

Park, H. Y., Lim, H., Yoon, Y. J., Follenzi, A., Nwokafor, C., Lopez-Jones, M., Meng, X., & Singer, R. H. (2014). Visualization of Dynamics of Single Endogenous mRNA Labeled in Live Mouse. Science, 343(6169), 422–424. 10.1126/science.1239200

Rao, V. R., Pintchovski, S. A., Chin, J., Peebles, C. L., Mitra, S., & Finkbeiner, S. (2006). AMPA receptors regulate transcription of the plasticity-related immediate-early gene Arc. Nature Neuroscience, 9(7), 887– 895. 10.1038/nn1708

Richter, J. D., Bassell, G. J., & Klann, E. (2015). Dysregulation and restoration of translational homeostasis in fragile X syndrome. Nature Reviews. Neuroscience, 16(10), 595–605. 10.1038/nrn4001

Roden, C., & Gladfelter, A. S. (2021). RNA contributions to the form and function of biomolecular condensates. Nature Reviews Molecular Cell Biology, 22(3), 183–195. 10.1038/s41580-020-0264-6

Sahoo, P. K., Lee, S. J., Jaiswal, P. B., Alber, S., Kar, A. N., Miller-Randolph, S., Taylor, E. E., Smith, T., Singh, B., Ho, T. S.-Y., Urisman, A., Chand, S., Pena, E. A., Burlingame, A. L., Woolf, C. J., Fainzilber, M., English, A. W., & Twiss, J. L. (2018). Axonal G3BP1 stress granule protein limits axonal mRNA translation and nerve regeneration. Nature Communications, 9, 3358. 10.1038/s41467-018-05647-x

Samadi, M., Hales, C. A., Lustberg, D. J., Farris, S., Ross, M. R., Zhao, M., Hepler, J. R., Harbin, N. H., Robinson, E. S. J., Banks, P. J., Bashir, Z. I., & Dudek, S. M. (2023). Mechanisms of mGluR-dependent plasticity in hippocampal area CA2. Hippocampus, 33(6), 730–744. 10.1002/hipo.23529

Sauerbeck, A. D., Gangolli, M., Reitz, S. J., Salyards, M. H., Kim, S. H., Hemingway, C., Gratuze, M., Makkapati, T., Kerschensteiner, M., Holtzman, D. M., Brody, D. L., & Kummer, T. T. (2020). SEQUIN multiscale imaging of mammalian central synapses reveals loss of synaptic connectivity resulting from diffuse traumatic brain injury. Neuron, 107(2), 257–273.e5. 10.1016/j.neuron.2020.04.012

Sawicka, K., Hale, C. R., Park, C. Y., Fak, J. J., Gresack, J. E., Van Driesche, S. J., Kang, J. J., Darnell, J. C., & Darnell, R. B. (2019). FMRP has a cell-type-specific role in CA1 pyramidal neurons to regulate autism-related transcripts and circadian memory. eLife, 8, e46919. 10.7554/eLife.46919

Sharangdhar, T., Sugimoto, Y., Heraud-Farlow, J., Fernández-Moya, S. M., Ehses, J., Ruiz de los Mozos, I., Ule, J., & Kiebler, M. A. (2017). A retained intron in the 3′-UTR of Calm3 mRNA mediates its Staufen2- and activity-dependent localization to neuronal dendrites. EMBO Reports, 18(10), 1762–1774. 10.15252/embr.201744334

Shiina, N., Shinkura, K., & Tokunaga, M. (2005). A Novel RNA-Binding Protein in Neuronal RNA Granules: Regulatory Machinery for Local Translation. The Journal of Neuroscience, 25(17), 4420–4434. 10.1523/JNEUROSCI.0382-05.2005

Sidorov, M. S., Auerbach, B. D., & Bear, M. F. (2013). Fragile X mental retardation protein and synaptic plasticity. Molecular Brain, 6, 15. 10.1186/1756-6606-6-15

Song, M. S., Moon, H. C., Jeon, J.-H., & Park, H. Y. (2018). Neuronal messenger ribonucleoprotein transport follows an aging Lévy walk. Nature Communications, 9, 344. 10.1038/s41467-017-02700-z

Steward, O., Bakker, C. E., Willems, P. J., & Oostra, B. A. (1998). No evidence for disruption of normal patterns of mRNA localization in dendrites or dendritic transport of recently synthesized mRNA in FMR1 knockout mice, a model for human fragile-X mental retardation syndrome. NeuroReport, 9(3), 477.

Trcek, T., Douglas, T. E., Grosch, M., Yin, Y., Eagle, W. V. I., Gavis, E. R., Shroff, H., Rothenberg, E., & Lehmann, R. (2020). Sequence-Independent Self-Assembly of Germ Granule mRNAs into Homotypic Clusters. Molecular Cell, 78(5), 941–950.e12. 10.1016/j.molcel.2020.05.008

Trcek, T., Grosch, M., York, A., Shroff, H., Lionnet, T., & Lehmann, R. (2015). Drosophila germ granules are structured and contain homotypic mRNA clusters. Nature Communications, 6(1), 7962. 10.1038/ncomms8962

Tübing, F., Vendra, G., Mikl, M., Macchi, P., Thomas, S., & Kiebler, M. A. (2010). Dendritically Localized Transcripts Are Sorted into Distinct Ribonucleoprotein Particles That Display Fast Directional Motility along Dendrites of Hippocampal Neurons. The Journal of Neuroscience, 30(11), 4160–4170. 10.1523/JNEUROSCI.3537-09.2010

Van Treeck, B., Protter, D. S. W., Matheny, T., Khong, A., Link, C. D., & Parker, R. (2018). RNA self-assembly contributes to stress granule formation and defining the stress granule transcriptome. Proceedings of the National Academy of Sciences of the United States of America, 115(11), 2734–2739. 10.1073/pnas.1800038115

Virtanen, P., Gommers, R., Oliphant, T. E., Haberland, M., Reddy, T., Cournapeau, D., Burovski, E., Peterson, P., Weckesser, W., Bright, J., van der Walt, S. J., Brett, M., Wilson, J., Millman, K. J., Mayorov, N., Nelson, A. R. J., Jones, E., Kern, R., Larson, E., … van Mulbregt, P. (2020). SciPy 1.0: Fundamental algorithms for scientific computing in Python. Nature Methods, 17(3), 261–272. 10.1038/s41592-019-0686-2

Wagle, S., Kraynyukova, N., Hafner, A.-S., & Tchumatchenko, T. (2023). Computational insights into mRNA and protein dynamics underlying synaptic plasticity rules. Molecular and Cellular Neuroscience, 125, 103846. 10.1016/j.mcn.2023.103846

Wang, G., Ang, C.-E., Fan, J., Wang, A., Moffitt, J. R., & Zhuang, X. (2020). Spatial organization of the transcriptome in individual neurons (p. 2020.12.07.414060). bioRxiv. 10.1101/2020.12.07.414060

Wang, H., Wu, L.-J., Zhang, F., & Zhuo, M. (2008). Roles of Calcium-Stimulated Adenylyl Cyclase and Calmodulin-Dependent Protein Kinase IV in the Regulation of FMRP by Group I Metabotropic Glutamate Receptors. Journal of Neuroscience, 28(17), 4385–4397. 10.1523/JNEUROSCI.0646-08.2008

Wang, H., & Zhuo, M. (2012). Group I Metabotropic Glutamate Receptor-Mediated Gene Transcription and Implications for Synaptic Plasticity and Diseases. Frontiers in Pharmacology, 3, 189. 10.3389/fphar.2012.00189

Wang, X., Zeng, W., Kim, M. S., Allen, P. B., Greengard, P., & Muallem, S. (2007). Spinophilin/neurabin reciprocally regulate signaling intensity by G protein-coupled receptors. The EMBO Journal, 26(11), 2768– 2776. 10.1038/sj.emboj.7601701

Wu, H., Zhou, J., Zhu, T., Cohen, I., & Dictenberg, J. (2020). A kinesin adapter directly mediates dendritic mRNA localization during neural development in mice. Journal of Biological Chemistry, 295(19), 6605– 6628. 10.1074/jbc.RA118.005616

Wu, L., Wells, D., Tay, J., Mendis, D., Abbott, M.-A., Barnitt, A., Quinlan, E., Heynen, A., Fallon, J. R., & Richter, J. D. (1998). CPEB-Mediated Cytoplasmic Polyadenylation and the Regulation of Experience-Dependent Translation of α-CaMKII mRNA at Synapses. Neuron, 21(5), 1129–1139. 10.1016/S0896-6273(00)80630-3

Zeitelhofer, M., Karra, D., Macchi, P., Tolino, M., Thomas, S., Schwarz, M., Kiebler, M., & Dahm, R. (2008). Dynamic interaction between P-bodies and transport ribonucleoprotein particles in dendrites of mature hippocampal neurons. The Journal of Neuroscience: The Official Journal of the Society for Neuroscience, 28(30), 7555–7562. 10.1523/JNEUROSCI.0104-08.2008

